# Barley shows reduced Fusarium Head Blight under drought and modular expression of differential expressed genes under combined stress

**DOI:** 10.1101/2023.02.15.528674

**Authors:** Felix Hoheneder, Christina E. Steidele, Maxim Messerer, Klaus Mayer, Nikolai Köhler, Christine Wurmser, Michael Heß, Michael Gigl, Corinna Dawid, Remco Stam, Ralph Hückelhoven

## Abstract

Plants often face simultaneous abiotic and biotic stress conditions. However, physiological and transcriptional responses of plants under combined stress situations are little understood. Spring barley is susceptible to Fusarium Head Blight (FHB), which is strongly affected by weather conditions. We therefore studied the potential influence of drought on FHB severity and responses in three differently susceptible spring barley varieties and found strongly reduced FHB severity in susceptible varieties under drought. Quantity of differentially expressed genes (DEGs) and strength of transcriptomic regulation reflected the concentration of physiological stress markers such as abscisic acid or fungal DNA contents. Infection-related gene expression associated rather with susceptibility than resistance. Weighted gene correlation network analysis uncovered 18 modules of co-expressed genes, which reflect the pathogen or drought response in the used varieties. A generally infection-related module contained co-expressed genes for defence, programmed cell death and mycotoxin-detoxification indicating that diverse genotypes use a similar defence strategy towards FHB albeit with different success. Further DEGs showed co-expression in drought or genotype-associated modules correlating with measured phytohormones or the osmolyte proline. The combination of drought stress with infection lead to highest numbers of DEGs and provoked a modular composition of single stress responses rather than a specific transcriptional readout.

**Highlight:** Co-expression network analysis reveals association of physiological stress markers and gene expression modules under biotic and abiotic stress

## Introduction

Under natural conditions, plants are facing multiple abiotic and biotic stress factors affecting growth, crop yield andbathogen defence. The simultaneous occurrence of two or more stress factors triggers complex regulatory responses resulting in synergistic, antagonistic or neutral effects on expression and balancing of stress responses (Choi *et al*. 2013; Zhang and Sonnewald 2017; Pandey *et al*. 2015; Ramegowda and Senthil-Kumar 2015; Pandey and Senthil-Kumar 2019). Many stress responses share common factors of plant signalling and metabolism, and co-occurrence of stressors is often fatal for crop productivity (Farooq *et al*. 2009; Suzuki *et al*. 2014; Zandalinas *et al*. 2021). Transcriptional analyses in *Arabidopsis thaliana* indicate that plants respond to combined abiotic and biotic stress with tailored regulatory responses differing from responses towards single stressors (Gupta *et al*. 2016). Stress-stress interactions are often complex and not well understood (Atkinson and Urwin 2012). One of the most limiting factors of plant growth, performance and crop productivity is water availability (Kang *et al*. 2009). In conjunction with global warming, strong and frequent drought periods are predicted to increase in near future and represent serious threats to food security and supply (Meza *et al*. 2020) and especially spring barley will strongly be affected by droughts (Olesen *et al*. 2011; Xie *et al*. 2018). A recent study further predicts that on a global scale proportions of fungal soil borne pathogens will increase with global warming (Delgado-Baquerizo *et al*. 2020). Hence, vulnerability of agriculture may rise with a simultaneously predicted increase in global food demands leading to increasing gaps in food security. In this context, understanding plant physiology under complex stress combinations could unfold regulatory pathways and marker genes important for advanced genotype selection and breeding of disease resistant and stress tolerant crops (Pandey and Senthil-Kumar 2019).

Fusarium head blight (FHB) is one of the most destructive diseases of small grain cereal crops worldwide and its epidemiology is strongly associated with weather conditions. The disease is caused by a pathogen complex of several *Fusarium* species such as *F. graminearum* and *F. culmorum*. Infections with *Fusarium* pathogens can cause yield loss but also quality losses and contamination with hazardous mycotoxins like deoxynivalenol (DON) or zearalenone (Wegulo *et al*. 2015). The occurrence and strength of the disease is dependent on multiple environmental factors especially temperature, air humidity and rainfalls around anthesis, which collectively influence disease incidence and severity in the field (Birr *et al*. 2020; Hoheneder *et al*. 2022). FHB resistance of wheat (Buerstmayr *et al*. 2021; Mesterhazy 2020; Buerstmayr and Lemmens 2015) and barley (Buerstmayr and Lemmens 2015; Ogrodowicz *et al*. 2020) is a quantitative trait expressed from various QTLs and hence difficult to exploit. Furthermore, effective chemical control of FHB is limited to a short time window at around anthesis, when primary head infection usually occurs. Additionally, control measurements and pathogenicity of the fungus strongly interfere with weather conditions, both increasing complexity of effective FHB management (Wegulo *et al*. 2015).

In order to obtain physiological markers for FHB responses under complex environmental conditions, understanding gene regulation networks in response to single and combined abiotic and biotic stressors is crucial. The transcriptional response of wheat and barley to FHB is increasingly well understood (Hameed *et al*. 2022; Kazan and Gardiner 2018) but little information is available for complex stress situations. Hormones regulate various stress responses and physiological and developmental processes in plants. Under drought, plant hormones act as notable endogenous plant growth regulators and mediate abiotic stress tolerance (Farooq *et al*. 2009) and pathogen defence (Bari and Jones 2009). Predominantly, abscisic acid (ABA) governs growth, root-shoot ratio and regulates stomatal conductance to reduce transpiration and water loss under drought over relatively long distances from the root to the shoot. However, while ABA improves abiotic stress tolerance during vegetative growth, the downstream effects of ABA during reproductive stage can be contrary (Dolferus *et al*. 2011). There is evidence that reproductive tissue is most sensitive to water deficit in general (Blum 2009) and drought stress strongly delays flowering or even leads to abortion of spikelets in cereal crops (Barnabás *et al*. 2008). When it comes to pathogen infection, the genetic regulatory mechanisms to balance hormonal regulations associated with specific stress responses are not sufficiently investigated, especially when invading microbes are able to manipulate regulatory networks in the host by biosynthesis of phytohormones (Bari and Jones 2009), which was previously described for *Fusarium* species (Dörffling and Petersen 1984; Jaroszuk-Ściseł*et al*. 2014; Luo *et al*. 2016; Qi *et al*. 2016) Salicylic acid (SA), jasmonic acid (JA) and ethylene are mainly involved in diverse pathogen defence responses against various pathogens and expression of pathogenesis related (PR) genes (Bari and Jones 2009). ABA induces several transcription factors, which contribute to specific abiotic stress but also pathogen defence regulations (Yao *et al*. 2021). SA is further associated as signal for systemic acquired resistance mechanisms. However, the function of hormonal defence signalling network is dependent on the pathogen’
ss trophic lifestyles (Adie *et al*. 2007), which is often not clearly classifiable and changes during pathogenesis of the hemibiotrophic *Fusarium* species from biotrophic to necrotrophic (Brown *et al*. 2010). However, the antagonistically acting SA and JA/ethylene associated pathways are both involved in responses to *Fusarium* infections (Wang *et al*. 2018). In particular, when it comes to combined stress, functions of either synergistically or antagonistically interacting plant hormones provoke complex stress responses. Hence, the host plant may prioritize particular stress responses, which can lead to either resistance or susceptibility (Bari and Jones 2009; Wang *et al*. 2018; Gupta *et al*. 2020). Although, previous studies revealed associations of ethylene (Chen *et al*. 2009; Xiao *et al*. 2013) auxin (Brauer *et al*. 2019) or ABA (Buhrow *et al*. 2021; Qi *et al*. 2016) with susceptibility or SA (Makandar *et al*. 2012), JA (Sun *et al*. 2016) and gibberellic acid (Buhrow *et al*. 2021) with resistance against *Fusarium* spp., roles of phytohormones remain complex and ambiguous. It was further shown, that their role during infection with *F. graminearum* strongly depends on distinct stages of infection demonstrating strong interactions of the pathogen with plant physiology (Ding *et al*. 2011; Ameye *et al*. 2015; Makandar *et al*. 2010; Makandar *et al*. 2012). Although, several biological and physiological mechanisms of *Fusarium* spp.-barley pathosystem were described, understanding of FHB susceptibility and resistance under complex environmental stress conditions poses new challenges but could support future breeding and selection of pathogen resistant and stress tolerant genotypes. For this purpose, the present study aims to investigate genotype-dependent quantitative resistance and regulatory networks of barley under infection with *Fusarium culmorum* in combination with drought stress. This revealed genotype-independent stress response markers, infection success-related gene expression clusters and a modular gene expression network under combined stress.

## Material and methods

### Description of greenhouse experiments

Three spring barley genotypes (Barke, Morex and Palmella Blue) were grown under controlled conditions in glass house cabins with air conditioning for temperature control. The genotypes were preselected according to different resistance to FHB as assessed in inoculation trials in the field (Hoheneder *et al*. 2022) and preliminary greenhouse.

Six barley grains were sown in each pot containing 3 L peat substrate (Einheitserde C700, Stender, Germany). 12 pots were prepared per barley genotype and stress treatment. We applied daily automatic watering and additional lightning for 16 h per day. Plants were randomized twice a week. Temperature was set to 18 °C (day) and 16 °C (night) with a relative air humidity of 60%.

Drought conditions were set for half of the pots from seven days before expected flowering (growth stage 65) on by a stop of automatic watering on separate flooding tables. To prevent plants from early and premature plant death due to fast desiccation of the substrate, each pot received little amount of water (50 mL) in the first three days after stopping daily irrigation. All plants without irrigation showed reduced turgor pressure and partial loss of green leaf area indicating strong drought stress of the plants. Continuously watered plants showed normal growth and phenology.

Seven days after the beginning of drought conditions, spikes of irrigated or drought stressed plants were sprayed with spray flasks till run-off either with *F. culmorum* spore solution or mock solution to obtain four different treatment contrasts (WM: watered-mock, WFc: watered-infected, DM: drought-mock, DFc: drought-infected). To maintain optimal moist conditions for infection (99% relative air humidity), spikes were covered and sealed with transparent polythene bags for two days. Respective mock sprayed plants were similarly treated. Two and four days after inoculation, individual spikes were cut from each pot and flash frozen in liquid nitrogen for further DNA, RNA and metabolite extraction.

### Preparation of Fusarium culmorum inoculum

Fugal inoculum was cultured and propagated according to (Linkmeyer *et al*. 2013). Therefore, three isolates of *F. culmorum* (Fc002, Fc03, Fc06 – culture collection, Chair of Phytopathology, Technical University of Munich) known to strongly infect barley spikes and to produce DON (Linkmeyer *et al*. 2013; Hofer *et al*. 2016; Hoheneder *et al*. 2022) were combined in equal amounts. Spore solution was adjusted to 50,000 conidia per mL tap water and contained 1 mL/L Tween 80 to improve wetting of the spike tissue. The respective mock solution contained the same amount of Tween 80.

### Sample preparation, DNA and RNA extraction from immature spike tissue

Each individual sample of each biological triplicate per treatment variation was divided into two pieces using one part for RNA extraction. The remaining spike tissues was put together to a pooled sample of three spikes for DNA extraction. After separation, the spike samples were immediately ground in liquid nitrogen and stored at minus 70 °C. DNA extraction was carried out according to the protocol of (Fraaije *et al*. 1999) with minor modifications as described by (Hofer *et al*. 2016). DNA concentration was adjusted to 20 ng total DNA µL^-1^ with nuclease free water. RNA extraction was performed with Direct-zol RNA Miniprep Plus Kit (ZymoResearch, USA) according to manufacturer’s protocol.

### Quantification of fungal DNA

Fungal and barley DNA was determined with qPCR according to (Nicolaisen *et al*. 2009) using species specific primers and 10-fold dilution series of pure target DNA as standards. Non template controls only contained water. Quantitative PCR reactions were carried out using Takyon Low ROX SYBR 2X MasterMix blue dTTP (Eurogentec, Belgium) with a AriaMx real-time PCR system (Agilent Technologies, USA). DNA contents of each sample was determined in duplicates. Finally, fungal DNA was normalized with barley DNA in pg *F. culmorum* DNA ng^-1^ barley^-1^ DNA^-1^.

### Library preparation for Illumina HiSeq2500 sequencing

Preparation of libraries for 3’-RNA sequencing was carried out with Lexogen QuantSeq 3’-RNA-Seq Library Prep Kit (FWD) (Lexogen, Austria) for Illumina sequencing according to manufacturer’s protocol. Quantification of input RNA was conducted with Qubit Fluorometer 2.0 (Invitrogen, USA) and Qubit RNA BR (broad range) Assay Kit (Invitrogen, USA) according to manufacturer’s protocol in a range of 10 to 1200 ng total RNA. Quantification of library size was performed with Qubit Fluorometer 2.0 und Qubit DNA High Sensitivity Assay Kit (Invitrogen, USA). A check for distribution of mRNA fragment size of each library was performed with am Agilent 2100 Electrophoresis Bioanalyzer (Agilent Technologies, USA). Final quantification of each library was determined according to Illumina qPCR guide with a KAPA SYBR Fast Mastermix Low ROX (Peqlab, Germany) in a QuantStudio 5 real-time PCR system (Applied Biosystems, USA). Final libraries were normalized with elution buffer (Qiagen, Germany) to a final concentration to 2 nM and pooled with an equal amount of each sample library. Denaturation and dilution of libraries for HiSeq Clustering was performed according to protocol A of user guide (Illumina) with a concentration for clustering of 10 pM.

### llumina HiSeq2500 sequencing

Sequencing of libraries was carried out on a HiSeq2500 sequencing platform using HiSeq Rapid SR Cluster Kit v2 (Illumina, USA) and HiSeq Rapid SBS Kit v2 (50 Cycle) with run parameters set for multiplexed single-reads (read 1: 100 cycles) and single-indexed reads of 7 cycles. HiSeq Control Software 2.2.70 was used for sequencing. Image analysis and base calling was carried out with Real-Time Analysis (RTA) 1.18.66.4. Fastq-files were generated with CASAVA BCL2FASTQ Conversion Software v2.20. Raw data have been stored on NCBI-Gene Expression Omnibus with the accession number GSE223521.

### 3’-RNAseq trimming, mapping and read count

The quality of the raw 3’-RNAseq data was analysed with FastQC (http://www.bioinformatics.babraham.ac.uk/projects/fastqc). The trimming step was done with Trimmomatic (Bolger *et al*. 2014) using the parameters ILLUMINACLIP:Illumina-SE.fasta:2:30:10 LEADING:3 TRAILING:3 SLIDINGWINDOW:4:15 MINLEN:40. We used Hisat2 (Kim *et al*. 2015) with parameters -U --rna-strandness FR to map the trimmed data to the Morex v2 assembly (Monat *et al*. 2019). In order to add an artificial 3’UTR because of the 3’-RNAseq reads, we used a custom script to elongate the last exon either by 3 kb or until the next gene in the gff file of Morex v2. The counts of the generated bam files were assigned to genes using featureCounts (Liao *et al*. 2014) with the parameters -t gene -s 1 -M -O.

### CPM calculation

CPM (counts per million) were calculated using the *edgeR*-package (Robinson *et al*. 2010). The CPMs were calculated from the “DGEList” object with log = FALSE, normalized.lib.sizes = TRUE.

### Differential gene expression and cluster analysis

The differential gene expression analyses were carried out using the *edgeR*-package (Robinson *et al*. 2010). We kept all genes with a CPM equal or bigger than 15 in at least 10% of all samples. To increase the signal-to-noise ratio for the analyses, we pooled the two time points (48h and 96h post infection) for each sample and calculated the differential expressed genes with the following contrast formula: “((variety_condition_48h + variety_condition_96h)/2) - ((variety_watered-mock_48h + variety_watered-mock_96h)/2)” with variety := Barke, Morex, Palmella Blue and condition := watered-infected, drought-mock, drought-infected.

The cluster-analyses of the DEGs was done with the Multiple Expression Viewer (MeV, (Howe *et al*. 2011) using the implemented self-organizing tree algorithm (SOTA) with the default parameters. As input we used all differentially expressed genes with an FDR < 0.05 in at least one variety for one certain stress condition. For each analyses the log_2_ fold-change values over all samples for the respective gene sets were used.

### GO enrichment analysis

GO-enrichment analyses were performed on the differentially expressed genes using the *topGO*-R-package (Alexa and Rahnenfuehrer 2022). The GO-terms for each barley gene were downloaded from https://biit.cs.ut.ee/gprofiler/gost. As gene universe we have used all genes found in our RNAseq analyses that passed the threshold of having a CPM equal or bigger than 15 in at least 10% of all samples (35657 genes). The enrichment analyses for “molecular function” and “biological process” was done using the following settings: algorithm=“weight01”, statistic=“fisher”. Shown are the graphs for the 5 most significant nodes.

### Weighted gene correlation network analysis (WGCNA)

Co-expression analyses were performed using the R-package WGCNA (Langfelder and Horvath 2008, 2012). First we calculated the mean of the CPM values for the three replicates for each time point, variety, treatment and stress condition. We then selected the 12818 genes which showed in the differential gene expression analysis a sig. regulation with an FDR < 0.05 in at least one variety. The mean CPM values of those genes were normalized using variance stabilizing transformation from the *vsn* R-package (Huber *et al*. 2002). For the WGCNA analysis we have used the default settings with a soft power of 6 for the calculation of network adjacency of gene counts and the topological overlap matrix, clustering the genes into 18 modules plus 42 unassigned genes collected in the grey module.

For all the external traits (phytohormones, plant metabolites and fungal DNA) that we have correlated with our modules, we previously calculated the mean of the three replicates for each time point, variety, treatment and stress condition. The mean value was than subjected into the network analysis.

For the network analysis, the 25 edges between any two genes with the highest edge weight in each module have been selected and the corresponding network of those selected edges was drawn using the *igraph* R-package (Csardi and Nepusz 2006). For better visualization the networks were manually adjusted.

### Measurement of phytohormones and plant metabolites

Contents of phytohormones and secondary plant metabolites (abscisic acid, abscisic acid glucoside, phaseic acid, dihydrophaseic acid, auxin, salicylic acid, salicylic acid glucoside and proline) was carried out with mass spectrometry according to methods described in (Chaudhary *et al*. 2020) and (Abramov *et al*. 2021). For this purpose, 150 mg of grinded barley spike tissue as used for RNA extraction was used to determine contents of phytohormones and plant metabolites in relation to fresh weight.

## Results

### Global transcription analysis reflects variety-dependent susceptibility to FHB and enhanced resistance under drought

To study variety-dependent and -independent stress responses in barley and the effect of drought stress on FHB, we grew three diverse spring barley varieties in pots under controlled conditions in the greenhouse. The varieties Barke, Morex and Palmella Blue were selected based on differences in their resistance against FHB in preliminary greenhouse experiments and in inoculation trials in the field (Hoheneder *et al*. 2022) with Barke showing the most resistant phenotype and Palmella Blue being the most susceptible variety. Morex showed an intermediate resistance phenotype and represents the variety with the sequenced reference genome for the later analysis. Further genotype characteristics and agronomic traits for all three varieties can be found in the supplemental table T1.

Plants were either continuously irrigated (watered samples) or irrigation was stopped (drought samples) seven days before flowering (GS 57-59), a time period in which weather conditions are of pivotal importance for FHB pathogenesis in the field (Hoheneder *et al*, 2022). Inoculation or mock-treatment of the spikes was carried out with *Fusarium culmorum* (*Fc*) spore suspension on watered samples (watered-Fc, WFc or watered-mock, WM) or drought samples (drought-Fc, DFc or drought-mock, DM) at mid of flowering (GS 65) and samples were taken 48 and 96 hours post-infection (hpi) (Fig. 1A).

**Fig. 1.**
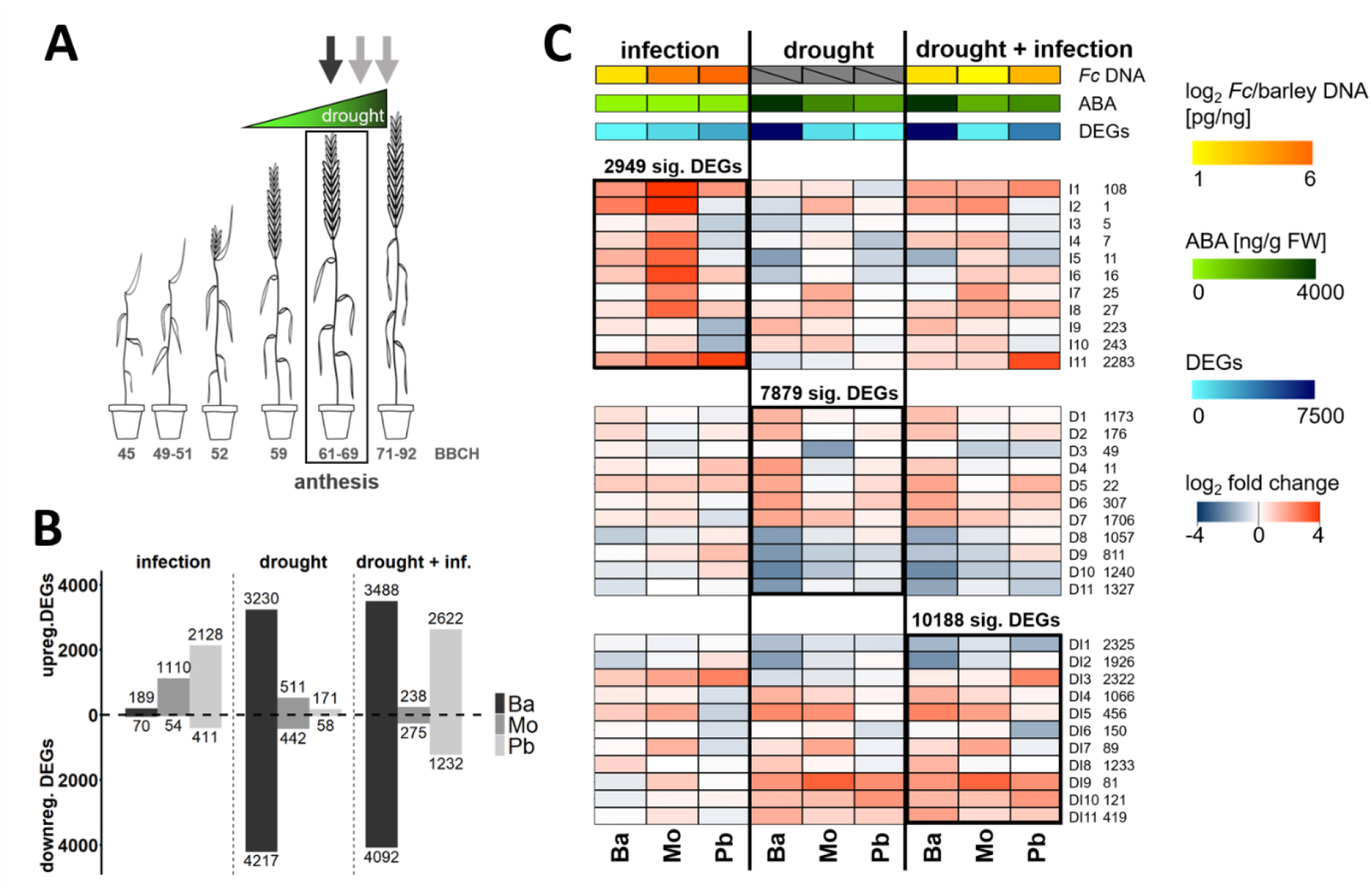
Global transcription analysis of variety-dependent, reduced susceptibility against FHB under drought stress in the greenhouse. (A) Experimental setup: Three different barley varieties (Barke, Ba; Morex, Mo; and Palmella Blue, Pb) were grown in pots under controlled conditions in the greenhouse. Drought stress was induced by stopped irrigation 7 days before anthesis (growth stage 61-69; dark grey arrow). Irrigated and drought-stressed plants were inoculated with conidia of *Fusarium culmorum* or mock (dark grey arrow) and samples were harvested 2 and 4 days post-inoculation (light grey arrows). (B) Number of DEGs. For each variety all treatments were compared against the watered, mock-infected samples. Differentially expressed genes were counted (FDR p<0.05) and number of DEGs were plotted. Upwards directed columns show the numbers of upregulated genes and downwards directed columns show downregulated genes. (C) Heat maps of key experimental outcomes and expression log_2_ fold changes of clusters of genes that behave similarly according to self-organizing tree algorithm. Amount of *Fusarium culmorum* DNA per ng barley DNA was determined by qPCR. Values were colour-coded with grey (0), yellow (log_2_ 1 pg *Fc* DNA ng^-1^ barley^-1^ DNA^-1^) to orange (log_2_ 6 pg *Fc* DNA ng^-1^ barley^-1^ DNA^-1^). The average DNA content from 2 and 4 dpi was calculated and the log_2_ used for data representation. Abscisic acid content [in pg ng^-1^ fresh^-1^ weight^-1^] in all spike samples was quantified using mass spectrometry. Values were colour-coded with light-green (0 ng ABA ng^-1^ fresh^-1^ weight^-1^) to dark-green (4000 ng ABA ng^-1^ fresh^-1^ weight^-1^). For each variety DEGs (relative to watered, not infected samples) with a FDR-corrected p-value < 0.05 were counted under each stress treatment and the quantity was colour-coded with light-blue (0) to dark-blue (7500). All genes which were significantly regulated (p < 0.05) in at least one variety in one particular stress treatment (infection: 2949 genes; drought: 7879 genes; drought + infection: 10188 genes) were subjected to a cluster analysis using a self-organizing tree algorithm (SOTA) using the expression pattern over all varieties and stress treatments. The respective stress treatment for which the DEGs were selected is highlighted by a black frame. All DEGs were split into 11 clusters, labelled with numbers 1-11 and a capital letter for the stress (I=infection, D=drought, DI=drought + infection) next to it and the corresponding number of genes in each cluster. The mean log_2_ fold change for each cluster was colour-coded ranging from blue (> −4) over white (0) to red (< 4).

In order to better understand the underlying differences in response to infection, drought stress or the combination of drought with infection, we analysed the gene expression of barley spikes using 3’-RNA-sequencing. Mapping of 3’-RNA reads on the reference genome, identified transcripts from 41746 (Barke), 42653 (Morex) or 41363 (Palmella Blue) barley genes in our samples. For 34969 genes, we obtained reliable reads from all three varieties. We compared for each variety all treatments against the watered, not-infected (watered-mock) samples and counted all significant (false discovery rate (FDR) corrected p < 0.05) differentially expressed genes (DEGs) (Fig. 1B). There were large variety-dependent differences in the amount of DEGs and in the distribution of up- and downregulated genes, ranging for example from 229 DEGs in Palmella Blue under drought stress over 953 genes in Morex up to 7447 genes in Barke. Under infection, by contrast, susceptible Palmella Blue showed most DEGs. In total and over all genotypes, we found 2949 DEGs after *F. culmorum* infection, 7879 DEGs under drought, and 10188 DEGs under drought plus infection. Because the gene sets overlap, this corresponds to a sum of 12818 DEGs (Supplemental data D1.1).

We used quantification of fungal DNA to assess the severity of infection in the same experiment. Under irrigation we observed strong differences in the amount of fungal DNA from Barke (Ba, 6 pg *Fc* DNA ng^-1^ barley^-1^ DNA^-1^) over Morex (Mo, 45 pg *Fc* DNA ng^-1^ barley^-1^ DNA ^-1^) to Palmella Blue (Pb, 70 pg *Fc* DNA ng^-1^ barley^-1^ DNA^-1^; averages of 48 and 96 h post inoculation) (Fig. 1C), corresponding well to the previously observed differences in basal resistance. When compared to this, all three varieties showed a reduced amount of fungal DNA when the plants were exposed to drought before infection (Ba 5 pg *Fc* DNA ng^-1^ barley^-1^ DNA^-1^; Mo 3 pg *Fc* DNA ng^-1^ barley^-1^ DNA^-1^; Pb 13 pg *Fc* DNA ng^-1^ barley^-1^ DNA^-1^).

We measured the accumulation of abscisic acid in all spike samples as one of the major responses to drought stress. We observed low values of abscisic acid in most of the irrigated samples, whereas the samples from drought-stressed barley showed clearly elevated levels of ABA, with strong genotype-dependent differences (Ba 4260 ng ABA g^-1^ FW^-1^; Mo 2057 ng ABA g^-1^ FW^-1^; Pb 1578 ng ABA g^-1^ FW^-1^; averages of 48 and 96 h post inoculation, 9 or 11 days after stop of irrigation) (Fig. 1C).

Despite strong differences in susceptibility to *F. culmorum* infection and in drought responses as measured by accumulation of ABA and the stress-associated amino acid proline (Supplemental data D2), our data also revealed DEGs that were regulated in all three genotypes under either infection-related stress (146 DEGs), drought stress (64 DEGs) or the combination of both (167 DEGs) (Supplemental data D1.2 - D1.4). Because our genotypes are diverse in geographic origin and pedigree (Supplemental table T1), those genes may serve as general variety-independent markers for the core response of barley to the respective stresses. In support of this, several of the DEGs in those lists are identical or homologous to previously reported DEGs in other barley or wheat genotypes infected by *Fusarium graminearum* or suffering from drought. Examples for such generally *Fusarium*-responsive genes are Fusarium resistant-orphan protein, tryptophan decarboxylases, anthranilate synthase, laccases, HvWRKY23 and DMR6-like 2-oxoglutarate and Fe(II)-dependent oxygenase genes (Buerstmayr *et al*. 2021; Low *et al*. 2020; Soni *et al*. 2020; Boddu *et al*. 2006; Boddu *et al*. 2007; Karre *et al*. 2019; Perochon *et al*. 2019; Tucker *et al*. 2021). Additionally, many enzyme genes that are potentially involved in detoxifying DON are among the commonly infection-regulated DEGs such as glycosyltransferases and Glutathione-S-transferase genes and a cysteine synthase (compare (Gardiner *et al*. 2010)). Prominent examples in that respect are HvUGT13248 (HORVU.MOREX.r2.5HG0384710) and HvUGT6 (HORVU.MOREX.r2.5HG0430540) (He *et al*. 2020; Michlmayr *et al*. 2018; Schweiger *et al*. 2010). In the list of genotype-independent drought-associated DEGs, we find previously reported dehydrins and late-embryogenesis-abundant protein genes, potential ABA-receptor complex PP2c protein phosphatases and a downregulated SAUR auxin response protein gene. 46 of the combined stress-associated variety-independent DEGs are also variety-independently regulated in one of the single stress situations. Those genes hence reliably showed the infection-related stress response even under drought and the drought-related response in additionally infected situations over all barley varieties (Supplemental data D1.5).

Interestingly, there are only two DEGs (HORVU.MOREX.r2.5HG0401150; HORVU.MOREX.r2.5HG0372030) that show significant regulation in all genotypes under combined stress but not after one of the single stresses in at least one genotype. Hence, barley seems to express no genotype-independent specific response to the applied combined stress.

The questions arose whether we find genotype-dependent stress markers in our data sets. We found many Barke-specific DEGs under drought and many Palmella Blue-specific DEGs after infection, because those genotypes showed the strongest gene expression responses to the respective stresses. We therefore more specifically asked whether there are Barke-specific DEGs under *Fusarium* stress that may explain higher quantitative resistance. This generated a list of 20 upregulated and 47 downregulated genes, which are neither differentially expressed in Morex nor in Palmella Blue (Supplemental data D1.6). Vice versa, we found 148 genes that were upregulated in both susceptible Morex and Palmella Blue but not in Barke (Supplemental data D1.7). Those lists possibly contain new factors for quantitative resistance or susceptibility to FHB.

With a self-organizing tree algorithm (SOTA) analysis, DEGs clustered according to their differential expression pattern over all varieties and treatments (Fig. 1C). Under infection with *F. culmorum*, the expression strength of infection-related cluster I11 (2283 genes; 77.4% of all infection-related DEGs) reflects the amount of fungal DNA (Fig. 1C). Cluster I9 and I10 (223 and 243 genes) show downregulation only in the most susceptible variety Palmella Blue. Taken together, 93.2% of all infection-related DEGs are in those clusters (I9-I11) and their expression strength rather reflects disease progression than resistance to FHB.

Under drought stress, the quantity of DEGs and the strength of expression in most of the clusters reflected the amount of ABA that we measured in individual varieties with Barke showing the strongest ABA accumulation (all drought-related clusters except D3, Fig. 1B, C).

The combination of drought stress with infection lead to the highest number of DEGs (10188 genes). Most of the clusters (DI1, 2, 4, 5, 9, 10, 11; 6394 genes, 62.8%) mirror very well the expression pattern under drought stress alone, indicating that those genes are mainly responsive to the drought stress. Cluster DI3 contains rather *Fusarium*-responsive genes, as they are not upregulated under drought stress alone, but show a strong infection-strength dependent regulation, similar to the before mentioned cluster I11. In particular, Palmella Blue, which is still strongly infected under drought, shows upregulation of genes in cluster DI3. Additionally, there are 3 clusters with variety-dependent expression that is similar in all stress situations (DI6 for Palmella Blue, DI7 for Morex and DI8 for Barke).

Taken together, we observed variety-dependent differences both in infection severity and strength of drought responses and all plants were less infected when plants were exposed to drought before infection compared to their respective controls. Each single stress provokes a mostly stress-specific gene expression response and the combination of drought stress and infection was mainly dominated by the firstly applied drought stress response in our experiment. However, in highly susceptible Palmella Blue many of combined stress-regulated DEGs are also infection-related DEGs. In many cases, number of DEGs and strength of gene regulation reflect an increase of a stress marker such as content of ABA or fungal DNA in the same samples.

### Expression network analysis supports global similarities in the FHB-responses, but also differences in drought and combined stress responses

For further analysis of the DEGs we performed a weighted gene co-expression network analysis (Zhang and Horvath 2005) to find DEGs that cluster into modules according to their expression pattern over all samples, time points and treatments (Fig. 2A), resulting in 19 modules of DEGs. 42 DEGs did not match any co-expression pattern (grey module) (Fig. 2A). The rest of the DEGs clustered into one of the remaining 18 modules, range in size from 63 genes up to 5287 genes in the turquoise module.

**Fig. 2.**
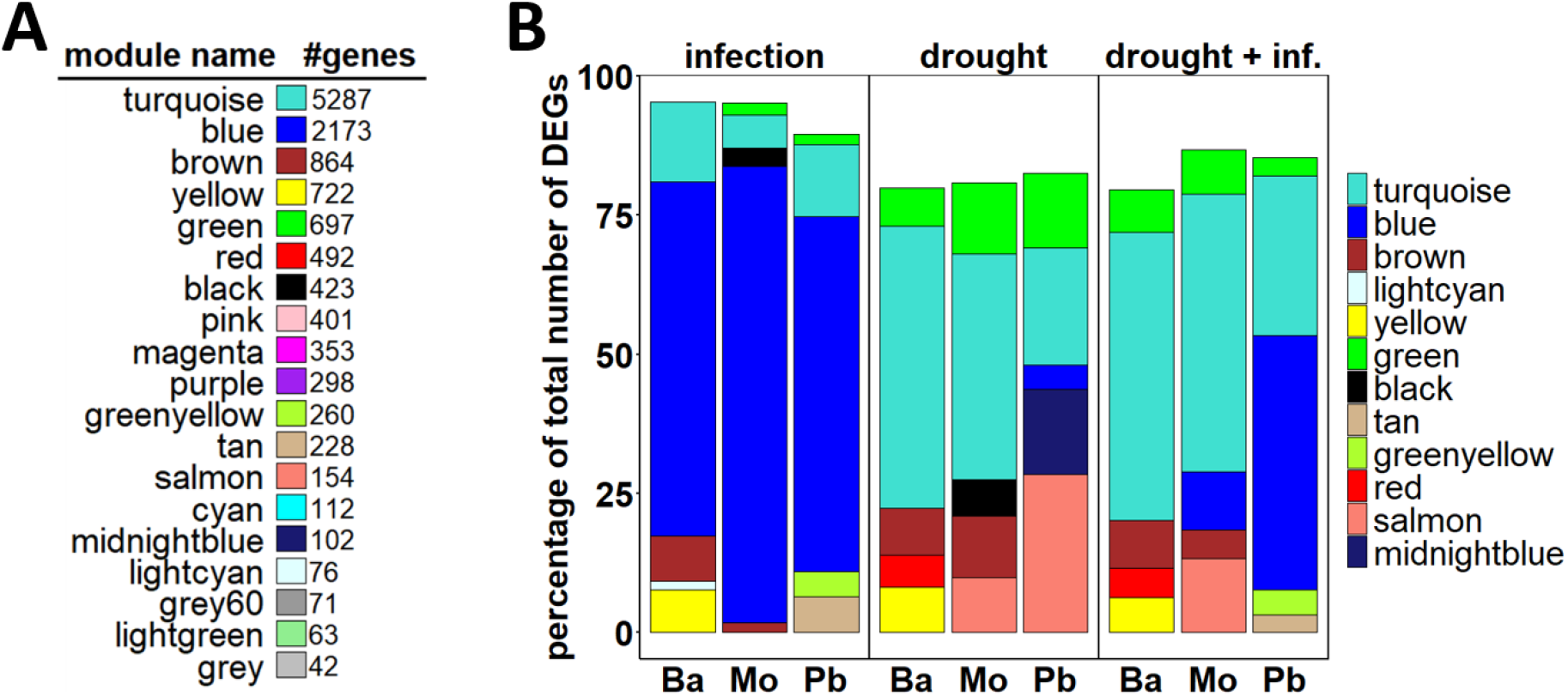
Modules of DEGs after weighted gene co-expression network analysis. (A) Co-expression modules. Based on the expression pattern over all time points, varieties and treatments all DEGs clustered into 18 modules (color-coded) with 63 to 5287 genes after WGCNA-analysis. 42 genes didn’t follow any co-expression pattern and were clustered in the grey module. The numbers reflect the amount of DEGs in each module. (B) Top 5 biggest co-expression modules in each variety for each stress treatment. We counted the DEGs in each module for each variety under each stress treatment. We show the 5 modules with most DEGs in each stress scenario in percentage relative to the total number of DEGs per variety (Barke, Ba; Morex, Mo; and Palmella Blue, Pb).

We checked for each barley variety and stress the relative contribution of modules to the overall stress response and displayed the five modules with the highest contribution to the sum of DEGs (Fig. 2B, supplementary Fig. S1). The blue module contains more than 60% of the infection-related largely upregulated DEGs for all three varieties. This may indicate that all three varieties show a similar response to infection with *F. culmorum* despite the fact that p-value based numbers of DEGs strongly differed. Gene ontology analyses ranked protein phosphorylation and response to biotic stress as most significantly enriched biological process (supplementary data D3 and D4A) and heme binding, DNA-binding transcription factor activity and protein serine/threonine kinase activity as most significantly enriched molecular functions in the blue module (supplementary data D3 and D4) possibly reflecting a general pathogen response in the blue module.

The drought stress response in Barke and Morex is dominated by the turquoise co-expression module but also the green module contributes to the drought response in all genotypes. In Palmella Blue the salmon module contains most DEGs but the turquoise module is also represented. Gene ontology analyses identified photosynthesis related functions as most significantly enriched in the turquoise module, and most of the genes are downregulated under drought (supplementary data D1.3, D3 and D4). The green module contains many similarly up-regulated genes and is most significantly enriched in gene ontology terms photorespiration, response to stress ending in defence response and embryo development ending in seed dormancy, and the salmon module with protein dephosphorylation and response to water (supplementary data D3 and D4).

After the combination of drought stress and infection most of the DEGs in Palmella Blue cluster in the blue module as seen for infection alone. In Barke and Morex the turquoise module is still the biggest one. Interestingly, none of the modules showed a significant association with the trait “combined stress” (see “DFc against all” in supplemental Figure S2A).

Additionally, we can also see genotype-related modules such as the yellow module being regulated in Barke under all three stress conditions but contributing less to the sum of DEGs in Morex and Palmella Blue DEGs (Fig. 2B, supplemental Figure S1).

The contribution pattern of individual modules to the overall stress response supports the previous observation that the combination of drought stress with infection is mainly dominated by the drought stress in Barke, because the distribution of the biggest modules is almost not altered in Barke under drought stress alone and under drought plus infection. In comparison, the distribution pattern of DEGs under drought plus infection matches better to infection alone in Palmella Blue (Fig. 2B, supplemental Figure S1). Morex shows a more complex response, but the drought stress pattern is largely recovered under combined stress.

### Co-expression modules correlate with FHB-severity, phytohormones or stress marker

For plant stress responses, little information exists on association of gene co-expression clusters with quantitative physiological traits of stress responses. We analysed module-trait associations by correlating WGCNA-module-sample eigengenes with determined contents of phytohormones and their derivates, abundance of the stress marker proline and the amount of fungal DNA to identify significant associations (Fig. 3).

**Fig. 3.**
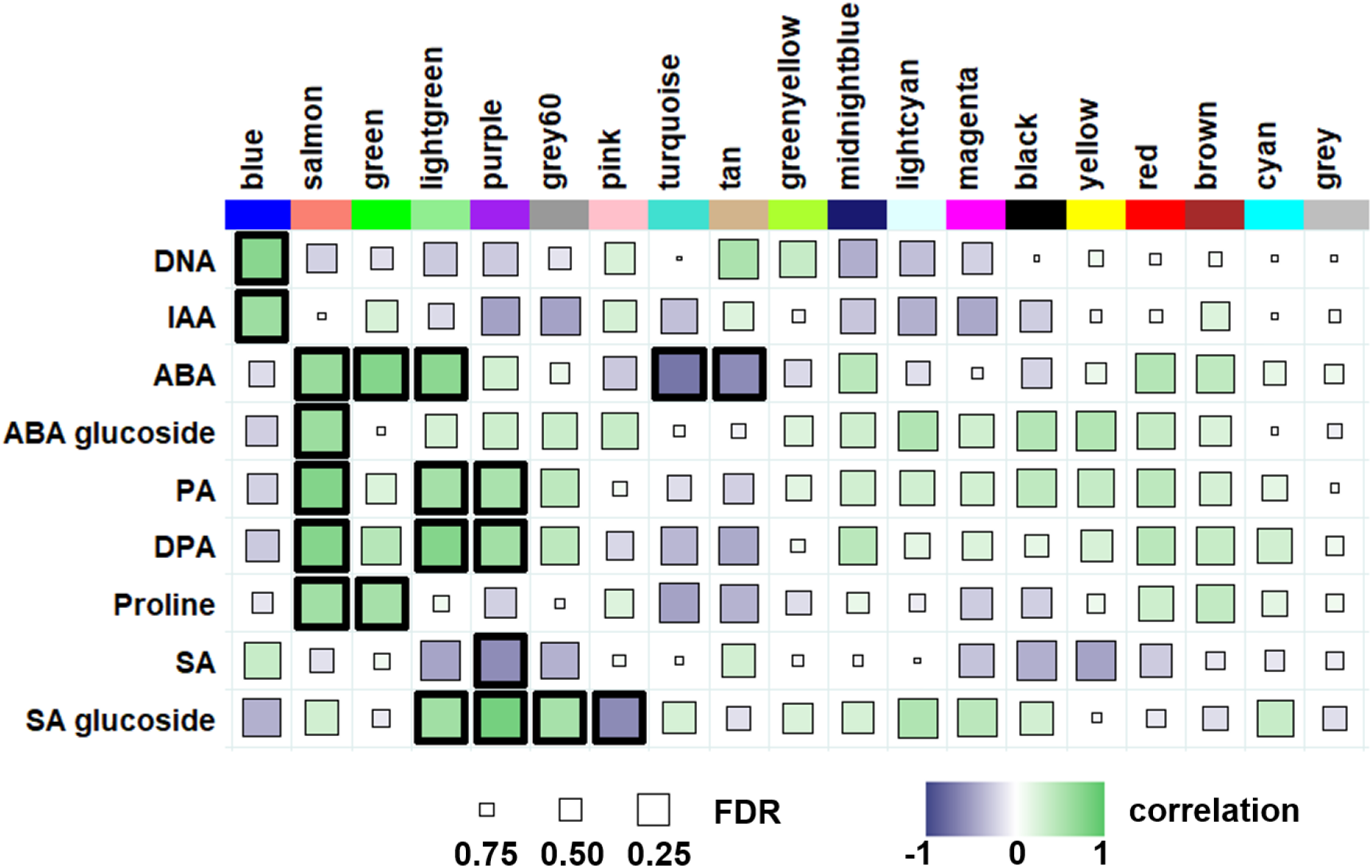
Relationships of consensus module eigengenes with FHB-severity, different phytohormones or stress markers. Each column in the table corresponds to a module and each row to one of the physiological traits: FHB-severity, a phytohormone, its derivate or proline as a stress marker. Module names are shown on top. Square colours in the figure represent the correlations of corresponding module eigengenes and measured stress parameters. The FDR-corrected p-values are coded by size (the bigger the square, the more significant). Highly significant correlations (p < 0.01) are highlighted with bold frames. DNA, *F. culmorum* DNA relative to plant DNA; ABA, abscisic acid; PA, phaseic acid; DPA, dihydrophaseic acid; IAA, indol 3-acetic acid; SA, salicylic acid.

The blue module is the only module which has a significant positive correlation with the amount of fungal DNA and the phytohormone auxin (IAA). The turquoise module shows the strongest negative correlation with ABA; most genes in this module are downregulated under drought. The green module has the highest positive correlation with the content of ABA. Similarly, to the turquoise module, the tan module shows a significant negative correlation with ABA contents. The salmon module, containing most of the DEGs in Palmella Blue under drought, shows a significant positive correlation with proline, ABA and the ABA derivates phaseic acid, dihydrophaseic acid and ABA glucoside. Hence, the module indicates a specific set of DEGs for stress regulation under drought. Additionally, many modules show by-trend contrasting correlations with salicylic acid or auxin versus ABA or derivates of ABA. The purple module is even significantly positively associated with phaseic acid and dihydrophaseic acid but negatively with salicylic acid. Hence, module-trait associations added functional value in showing significant linkages between stress physiology and complex gene expression patterns (Fig. 3).

### Co-expression levels reveal potential hub genes in selected modules

For a better understanding of the inter-connection and functionality-related linkage between the most co-regulated genes within selected modules we took the 25 edges with the highest edge weight from each of those modules. The edge weight represents the connection strength between two nodes, corresponding to the co-expression of those two genes.

The blue module is the only one with a significant positive correlation with the amount of fungal DNA within barley spikes (Fig. 3) and most of the heavy edges are directly connected with the DON-detoxifying Glycosyltransferase HvUGT13248 (HORVU.MOREX.r2.5HG0384710) (Mandalà *et al*. 2019; Schweiger *et al*. 2010) (Fig. 4A, B, supplemental table D5). All of the selected genes are infection-specifically expressed and the expression strength partially reflects the amount of fungal DNA detected in each variety, with Palmella Blue having the highest amount of fungal DNA detected by qPCR (see Fig. 1C). Most of the genes show a positive fold change after infection of watered and drought stressed plants, but no significant regulation under drought stress alone (Fig. 4B). Manual gene reannotations suggested a possible function of genes in DON-toxin response (HvUGT13248 and a GSTU6-like Glutathione S-transferase, HORVU.MOREX.r2.7HG0532730), plant defence and regulation of cell death (e.g. heat shock transcription factor similar to rice lesion mimic gene *spl7*, HORVU.MOREX.r2.1HG0067240; a NAC transcription factor, similar to OsNAC4 involved in cell death responses, HORVU.MOREX.r2.3HG0247070; disease resistance protein a, which represents a Toll-interleukin receptor 1 domain only protein, HORVU.MOREX.r2.2HG0110590) (Kaneda *et al*. 2009; Yamanouchi *et al*. 2002) and of plant immunity (homologs of Arabidopsis PUB21 and PUB23 plant U-box domain-containing family proteins, HORVU.MOREX.r2.6HG0521570, HORVU.MOREX.r2.7HG0555030) (Trujillo *et al*. 2008).

**Fig. 4.**
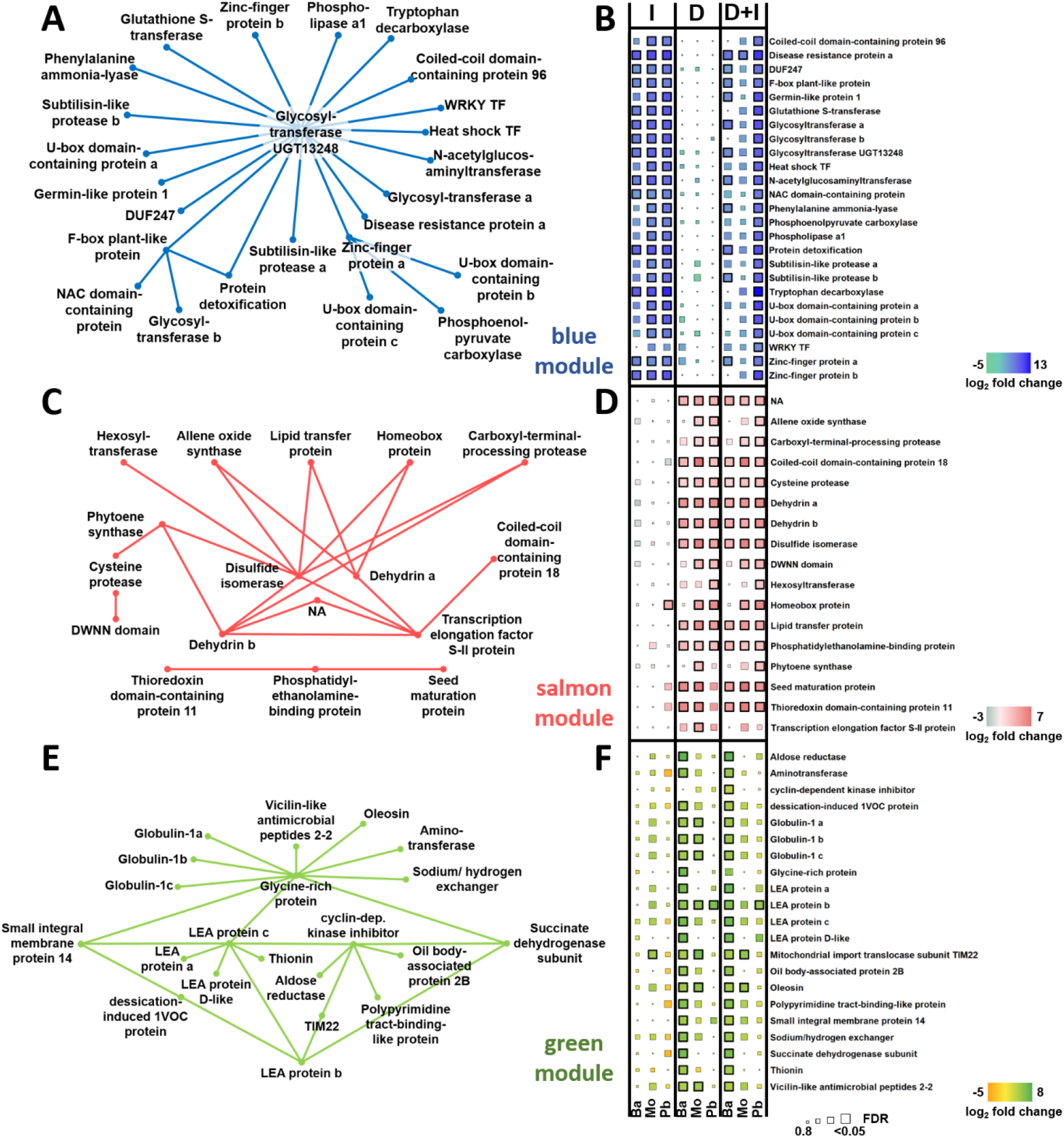
Co-expression levels reveal potential hub genes in selected modules. For three co-expression modules (blue, salmon and green), the 25 edges with the highest edge weight have been selected and their corresponding network was schematically drawn (A blue module, C salmon module, E green module). Each node represents one gene, and in the heat maps on the left the log fold changes of the gene expression values are shown with the respective colour codes and individual scales for each module (B blue module, D salmon module, F green module). The size of the square represents the false discovery rate-corrected p-values (FDR), with bigger squares indicating higher confidence levels. The bold squares indicate a significant FDR p-value smaller than 0.05. For gene identifiers, see supplemental dataset D5.

The salmon module shows positive correlation with drought stress associated ABA and ABA-derivates and the drought stress marker proline but no association with fungal infection success (Fig. 3). The genes corresponding to the 25 highest co-expression values are in all three varieties upregulated under drought stress, both alone and in combination with infection (Fig. 4D, supplemental table D5). Typical drought and dehydration stress-associated genes are among the strongly co-expressed genes. These are for instance dehydrins (HORVU.MOREX.r2.6HG0516710, HORVU.MOREX.r2.6HG0516720) or a seed maturation protein (HORVU.MOREX.r2.7HG0529950). Additionally, we find a phytoene synthase potentially involved in ABA biosynthesis (HORVU.MOREX.r2.5HG0419050) and hexosyltransferase similar to galactinol synthase 2, a drought stress tolerance protein that catalyses raffinose synthesis for osmoregulation (HORVU.MOREX.r2.2HG0090050) (Gu *et al*. 2016; Li *et al*. 2020; Selvaraj *et al*. 2017) and an ABA-related Arabidopsis AtHB-7-like homeobox transcription factor (HORVU.MOREX.r2.6HG0496860) (Söderman *et al*. 1996).

The green module is significantly correlated with ABA and proline but not fungal infection success (Fig. 3). The genes corresponding to the 25 highest co-expression values are, similar to what we found for the salmon module, genes associated with responses to dehydration stress, like late-embryogenesis abundant proteins (HORVU.MOREX.r2.1HG0049080, HORVU.MOREX.r2.1HG0049090, HORVU.MOREX.r2.3HG0213440, HORVU.MOREX.r2.5HG0352360), potential osmoregulatory sodium/hydrogen_exchanger (HORVU.MOREX.r2.7HG0560260) (Brini *et al*. 2007) or desiccation-induced 1VOC proteins (HORVU.MOREX.r2.4HG0322480) (Mulako *et al*. 2008) (Fig. 4E, supplemental table D5). A lot of the genes show a stronger regulation in Barke compared to the two others varieties potentially reflecting higher ABA contents in Barke under drought (Fig. 4F).

The turquoise module is by far the biggest module (5287 genes) (Fig. 2A) and is negatively correlated with ABA (Fig.3). Most of the strongest co-expressed genes are either associated with photosynthesis or are associated with translation (Suppl. Fig. S3A+B, supplemental table D5), two process being already known to be downregulated under stress.

The midnightblue module is not associated with any measured trait (Fig. 3), but is among the modules dominating the drought stress response of especially Palmella Blue (Fig. 2B). Interestingly, 8 out of the highly connected 11 genes with strongest co-expression are annotated as heat-shock proteins (Suppl. Fig. S3C+D, supplemental table D5).

Taken together the correlation of the modules and phytohormones or stress-markers is further strongly supported by the predicted functions of most strongly co-expressed genes in each module. Corresponding gene annotations support the hypothesis that the blue module may partly link to DON-responses and cell death during differentially progressing pathogenesis in the different varieties. The salmon and green module showed a significant correlation with several typical drought stress markers, like ABA or proline, and also the genes showing the strongest co-expression in these modules strongly support a key function in the response to drought stress and ABA.

## Discussion

Comparably little is known about the response of plants to a combination of different stresses and there is a high demand for understanding complex stress responses in crop plants (Rivero *et al*. 2022). Severity, frequency and combination of stresses collectively define the likelihood that stress results in tissue damage or complete organ or plant death (Buchanan 2000). Literature provides examples for plant responses to diverse simultaneous stresses that act additive or mutually inhibitory or synergistic (Ben Alaya *et al*. 2021; Loo *et al*. 2022; Rivero *et al*. 2022). We wanted to better understand how drought, which increasingly often occurs before flowering and seed set in mid European barley growing areas, may influence FHB infection. We used diverse spring barley genotypes that have different degrees of quantitative resistance to FHB under field and glasshouse conditions (Hoheneder *et al*. 2022) to survey a broad spectrum of possible responses in barley. We observed strong differences in the amount of the drought stress hormone ABA (Fig. 1C; Supplemental data D2) whereas the drought stress metabolite and osmolyte proline accumulated more uniformly among the three genotypes (Supplemental data D2). This might indicate that the genotypes experienced a similar change of water potential but Barke particularly strongly responded to this by accumulation of ABA and the strongest drought-related gene expression response. In contrast, Barke reacted least to *F. culmorum* infection, although it showed the highest degree of quantitative resistance as assessed by fungal DNA in infected ears (Fig. 1). By contrast, highly susceptible Palmella Blue showed most DEGs and strongest upregulation under infection. Morex showed an intermediate response in both aspects. Therefore, the global gene expression rather reflects a transcriptional response associated with successful FHB pathogenesis. However, many genes that are significantly regulated in susceptible barley are also infection-responsive in more resistant Barke albeit at a lower level and not significantly different from controls (Fig. 1; Supplemental data D1.1). We speculate that in less susceptible Barke less tissue gets in direct contact to the developing fungus and our low numbers of DEGs in most resistant Barke may be biased due to a dilution effect from healthy parts of the tissue. Nevertheless, high numbers and strong regulation of DEGs in susceptible barley do not reflect efficient defence, an observation that has been made before for FHB in wheat (Biselli *et al*. 2018; Buerstmayr *et al*. 2021; Wang *et al*. 2018).

Application of drought stress before FHB infection resulted in less successful pathogenesis. This effect was strong in Morex and moderate in Palmella Blue. In little susceptible Barke, drought did not have a major impact on FHB development. Accordingly, global gene expression responses to the fungus were reduced in susceptible barley under drought. However, in Palmella Blue, the amount of DEGs that appear to be primarily infection-responsive under combined stress was still high, and genes showed strong upregulation. Again, strong gene expression responses to fungal infection associate with fungal success, even if the success is limited under drought.

SOTA analysis allowed for a quick overview about global gene expression patterns. Generally, SOTA-resolved gene expression suggested that most combined stress regulated clusters showed a modular gene expression trend. This means that they often reflect gene expression that similarly appeared under drought or infection alone. We did not find clear indications for a specific response of barley to the combined stress but data rather support a combination of the drought and the infection response. If we consider the 2949 infection-related DEGs, we see that most clusters show a lower responsiveness under combined stress, when compared to infection stress alone. However, data do not resolve whether this reflects an inhibitory effect of drought stress on the pathogen response or rather less fungal infection success, although we maintained optimal infection conditions by covering barley spikes with polythene bags for two days after inoculation. Morex profited most strongly from drought when considering reduced fungal infection success. Interestingly, quite some clusters from the combined stress situation, show also mild pathogen- and drought-responsiveness in Morex (Fig. 1C, clusters DI5, 7, 9-11). It is tempting to speculate, that in this case the drought response positively added to the pathogen response and supported effective pathogen defence.

We used the full complexity of our data set and all DEGs to calculate gene co-expression networks by WGCNA. This uncovered modules of co-expressed genes, which largely reflect the pathogen response (blue module) or drought response (turquoise, green, salmon modules) in the used varieties. Similar information was extracted from the SOTA and as such not a surprise, e.g. infection related clusters of the SOTA analysis were rediscovered in the blue module of the WGCNA (supplementary dataset D1.1). Added value of the WGCNA derived from physiological trait association and diving into most strongly co-expressed gene sets (arbitrarily extracted 25 heaviest edges of big modules). Association of modules with physiological data of fungal development and of plant stress markers supported that the blue module is positively correlated with fungal DNA and auxin contents (Fig. 3). Previous reports from wheat FHB had already reported a connection between FHB susceptibility and auxin contents or responses (Brauer *et al*. 2019; Luo *et al*. 2016; Wang *et al*. 2018). This could hence reflect a general association in FHB of *Triticeae*. The blue module subnetwork of 25 most strongly co-regulated genes centered around the DON-detoxifying glycosyltransferase HvUGT13248 and more genes with predicted functions in toxin-response, dampening of immune-responses and programmed cell death regulation (Kaneda *et al*. 2009; Mandalà *et al*. 2019; Buerstmayr *et al*. 2021) (Fig. 4A). In this context it is interesting, that overexpression of the barley cell death suppressor BAX Inhibitor-1 enhances resistance to *F. graminearum* ear and crown infection in barley (Babaeizad *et al*. 2009). Since gene selection was based solely on edge weight in the network, and several discovered genes have partially described function in *Fusarium* spp. or general immune response, outcome suggests that our pipeline was suitable for finding potentially pivotal gene functions in big co-expression modules. This is supported by the fact that also the green and salmon module subnetworks of heavy edges reflect gene annotations known from drought responses and seed maturation (Fig. 4C, E). Self-pollinating barley is fertilized already before open flowering and seed set is initiated. We applied drought stress before open flowering and our data suggest that seed maturation was accelerated under drought, storage proteins and dehydrins are upregulated and also sugar osmolytes may enrich. This change in plant resource allocation could have also influenced fungal infection success because quick ripening might have limited the hemibiotrophic fungus in access to plant metabolites.

The interplay of biotic and abiotic stress responses is complex and difficult to study. Literature provides examples of mutually inhibitory or supportive stress responses, such that combined or consecutive stresses may result in enhanced or reduced stress resistance. Often plant hormone-triggered pathways act antagonistically in complex interactions with the environment, which may have evolved as a measure of checks and balances to fine tune responses for optimal fitness. Our data also show multiple tendencies of opposite correlation of gene expression in WGCNA modules and salicylic acid versus ABA or auxin versus ABA and related metabolites (compare grey-blue versus green squares in vertical columns of Fig. 3). Perhaps, ABA responses and early ripening limited pathogenesis-associated auxin responses and thereby reduced susceptibility to fungal infection.

The question arises, whether we can generalize the observations we made in terms of drought associated resistance to fungal infections in barley. Indeed, a recent study found many drought stress related genes to be differentially expressed during Fusarium crown rot development in barley and wheat (Su *et al*. 2021). In a similar context, a NAC transcription factor (HvSNAC1), involved in drought tolerance, was found to mediate resistance towards Ramularia leaf spot disease of barley (McGrann *et al*. 2015). We observed earlier, that Barke and fourteen other spring barley genotypes showed enhanced resistance to Ramularia leaf spots when artificially exposed to long lasting drought stress in the field. However, in open fields, severe Ramularia leaf spots occurred even in hot and dry summers if single short rain events provided sufficient leaf wetness (Hoheneder *et al*. 2021). It is hence difficult to generalize that drought would mitigate severity of fungal infections. The timing of drought stress application may further be pivotal for its influence on fungal infection success. In preliminary experiments, we found that drought can also enhance development of FHB disease, when applied simultaneously instead of before infection. It remains speculative, whether and how altered gene regulation is depending on the order of occurring abiotic and biotic stress situations. We found several NAC and WRKY transcription factors regulated under combined stress and clustering in the blue module (Supplemental data D1.1), which reveals importance of single and versatile acting genes on multiple stress responses. This and future studies may provide an increasing amount of data that allows for a deep understanding of how stress responses interact depending on plant genotype and phenology and timing and combination of stresses.

The complexity of gene expression data is a boon and a bane of combined stress response analyses. In front of thousands of stress-regulated genes in our study, using diverse genotypes allowed for identification of genotype-independent stress markers that are reliable both under single and combined stresses (Supplemental data D1). Computational association of gene expression modules with physiological or agronomic traits appears helpful for interpretation of complex data. Enhancing the complexity of experimental setups and data points also supports co-expression analyses and identification of subnetworks that may function in stress resistance or susceptibility and can identify trait-associated candidate genes for functional analysis.

## Supporting information

suppl.D1-DEGs

suppl.D2-Phytohormones_DNA

suppl.D3-GOterms-statistics

suppl.D4-GOterms-hierarchicalTrees

suppl.D5-moduleInformation-top25

suppl.FiguresS1-S3

suppl.T1-variety-information

## Supplementary data

*Table T1*. Detailed information about the three used barley varieties

*Figure S1*. Full list of co-expression modules in each variety for each stress treatment

*Figure S2*. Relationships of consensus module eigengenes with binary comparisons

*Figure S3*. Co-expression levels reveal potential hub genes in selected modules

*Supplemental data D1*. List of DEGs

*Supplemental data D2*. Data table with measured phytohormones, stress markers and fungal DNA

*Supplemental data D3*. Statistics of GO term enrichment for molecular function and biological process in selected modules

*Supplemental data D4*. Hierarchical trees of GO term enrichment for molecular function and biological process in selected modules

*Supplemental data D5*. Gene numbers, abbreviations and CPM values for genes with the highest co-expression values in selected module

## Abbreviations

ABA: abscisic acid
Ba: Barke
DFc: drought stress, infected with *Fusarium culmorum*
DEG: differently expressed gene
DM: drought stress, mock-inoculated
DPA: dihydrophaseic acid
*F*.: *Fusarium*
Fc: *Fusarium culmorum*
FHB: Fusarium Head Blight
IAA: indol 3-acetic acid
Mo: Morex
PA: phaseic acid
Pb: Palmella Blue
SA: salicylic acid
SOTA: self-organizing tree algorithm
WFc: watered, infected with *Fusarium culmorum*
WGCNA: weighted gene correlation network analysis
WM: watered, mock-inoculated
WRKY: WRKY transcription factor

## Acknowledgements

The authors thank Lena Forster and Carolin Hutter for technical assistance in the laboratory and greenhouse. We further thank the gardener and technician team of the Plant Technology Center at Technical University of Munich for maintenance of greenhouse experiments.

## Author contributions

FH, CES, MH and RH: conceptualization.

FH, CES and RH wrote the manuscript

FH and conducted the greenhouse experiments.

CW performed the RNA sequencing.

MG and CD measured the phytohormones and stress markers.

MM and KM mapped the RNAseq reads to Barley genome.

CES, NK, RS and RH analysed the obtained sequencing results.

