## Supplementary material for "Barley shows reduced Fusarium Head Blight under drought and modular expression of differential expressed genes under combined stress": suppl.D4-GOterms-hierarchicalTrees

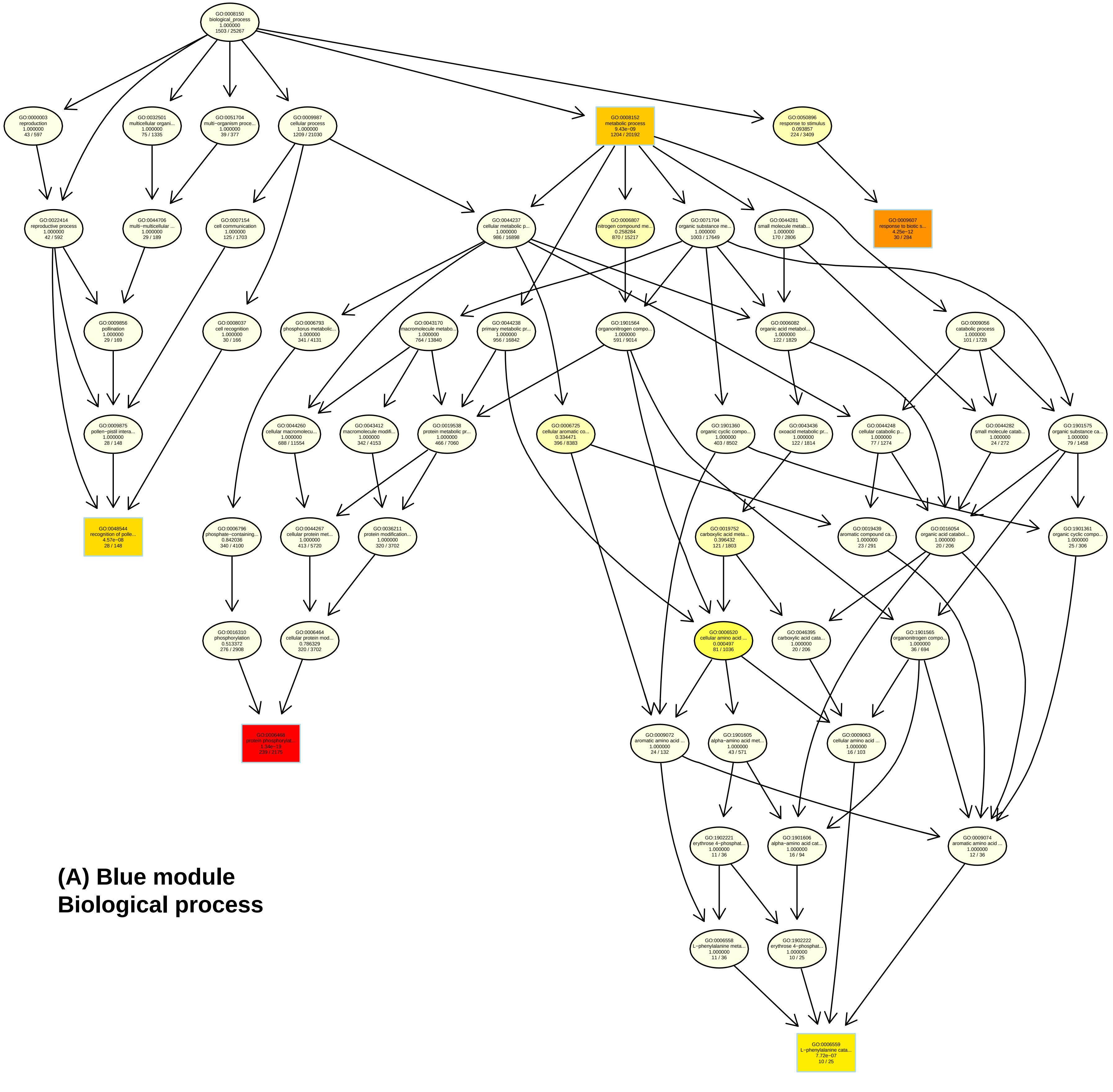


**(A) Blue module**
**Biological process**

### (B) Blue module
### Molecular function

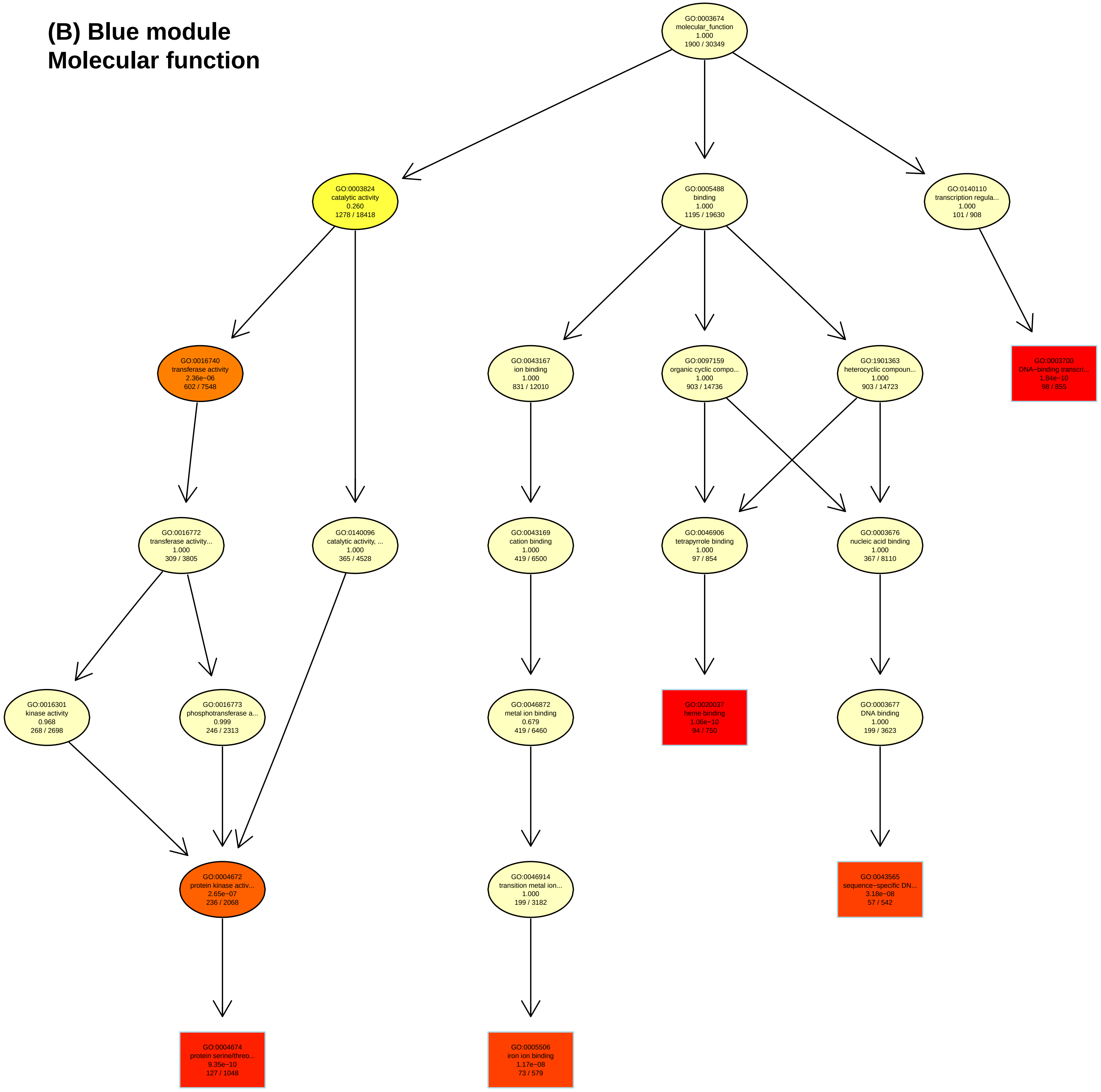

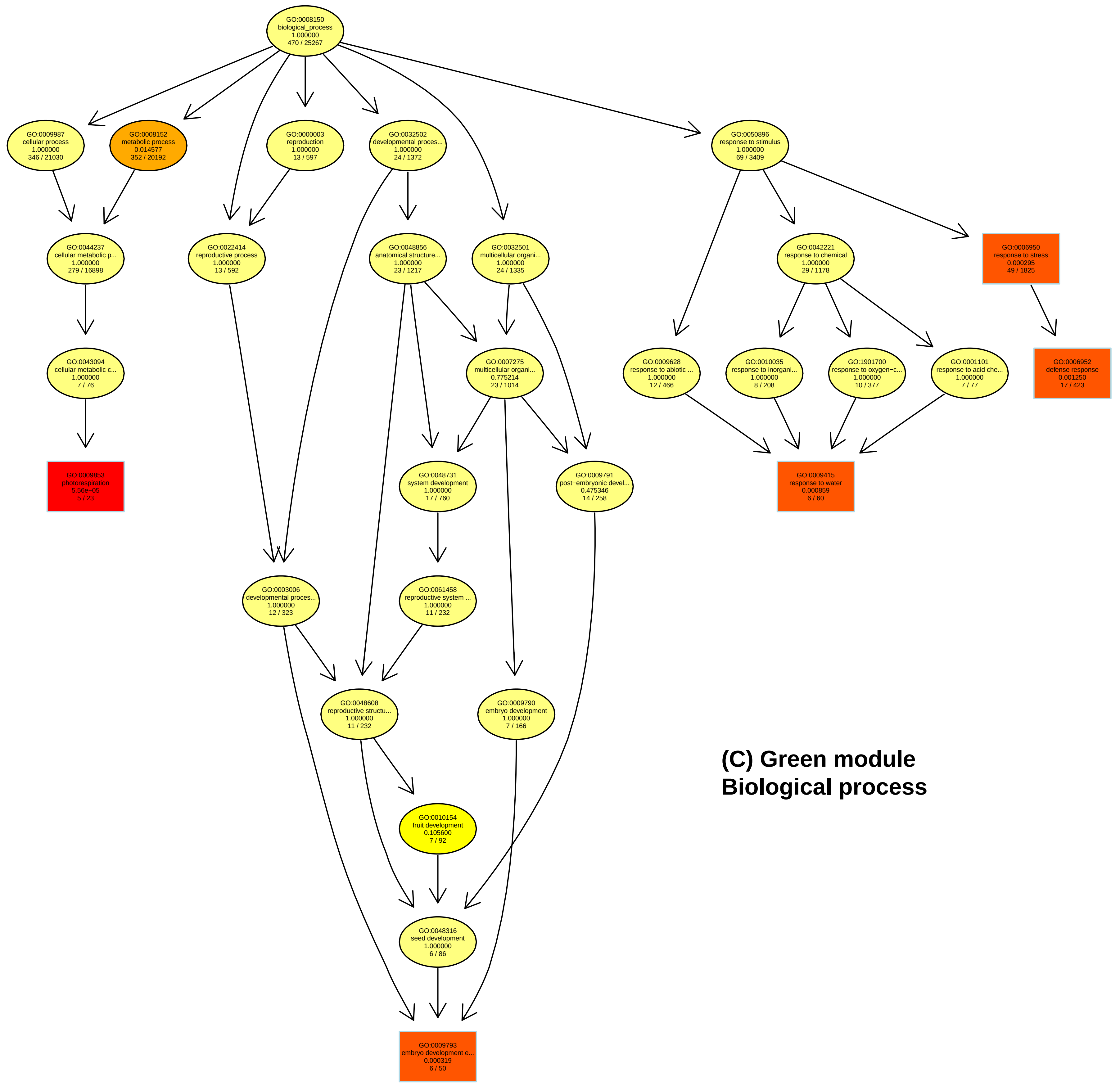


**(C) Green module**
**Biological process**

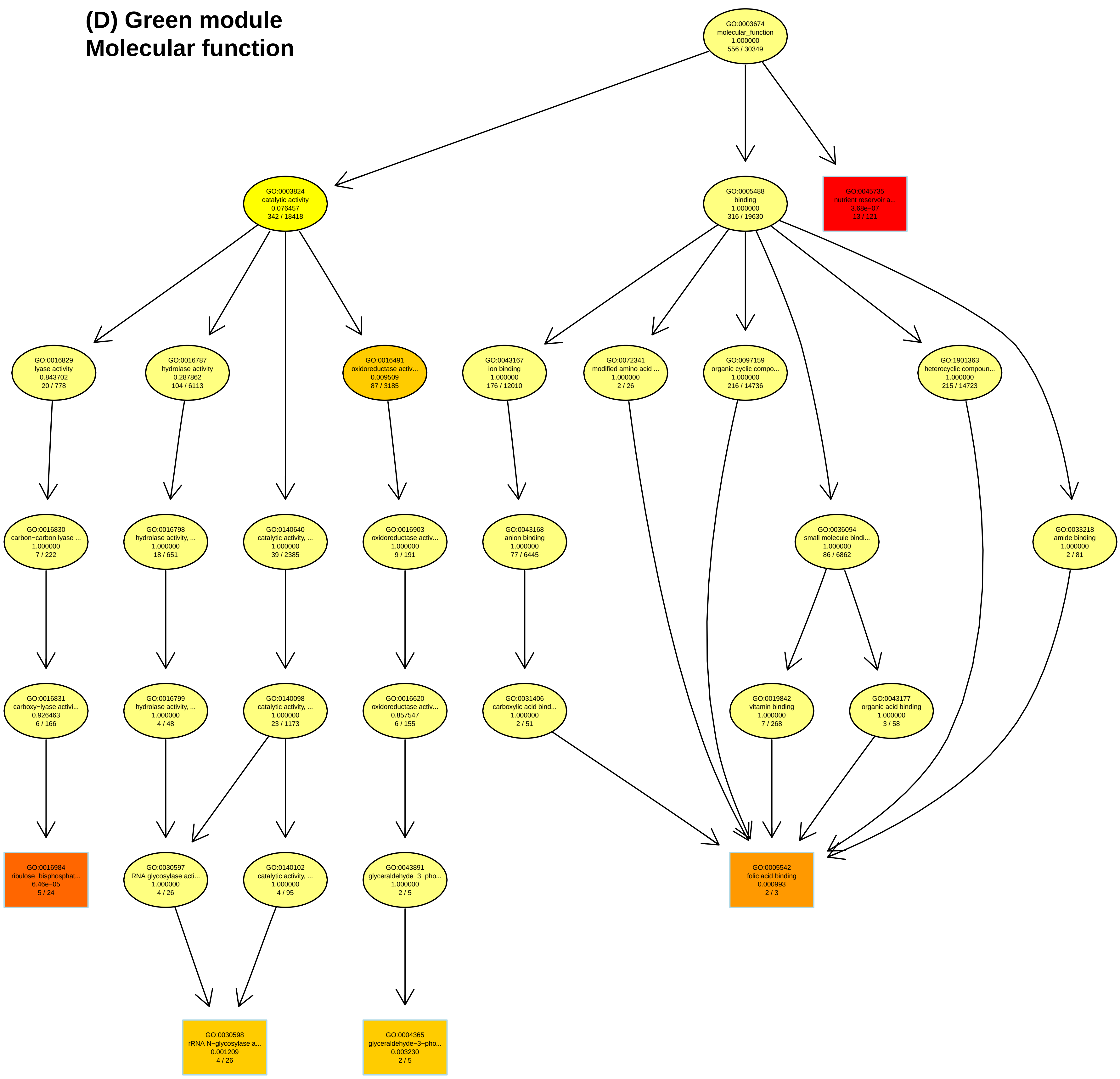
(D) Green module
Molecular function
GO:0003674
molecular_function
1.000000
556 / 30349
GO:0003824
catalytic activity
0.076457
342 / 18418
GO:0005488
binding
1.000000
316 / 19630
GO:0045735
nutrient reservoir a...
3.68e−07
13 / 121
GO:0016829
lyase activity
0.843702
20 / 778
GO:0016787
hydrolase activity
0.287862
104 / 6113
GO:0016491
oxidoreductase activ...
0.009509
87 / 3185
GO:0043167
ion binding
1.000000
176 / 12010
GO:0072341
modified amino acid ...
1.000000
2 / 26
GO:0097159
organic cyclic compo...
1.000000
216 / 14736
GO:1901363
heterocyclic compoun...
1.000000
215 / 14723
GO:0016830
carbon−carbon lyase ...
1.000000
7 / 222
GO:0016798
hydrolase activity, ...
1.000000
18 / 651
GO:0140640
catalytic activity, ...
1.000000
39 / 2385
GO:0016903
oxidoreductase activ...
1.000000
9 / 191
GO:0043168
anion binding
1.000000
77 / 6445
GO:0036094
small molecule bindi...
1.000000
86 / 6862
GO:0033218
amide binding
1.000000
2 / 81
GO:0016831
carboxy−lyase activi...
0.926463
6 / 166
GO:0016799
hydrolase activity, ...
1.000000
4 / 48
GO:0140098
catalytic activity, ...
1.000000
23 / 1173
GO:0016620
oxidoreductase activ...
0.857547
6 / 155
GO:0031406
carboxylic acid bind...
1.000000
2 / 51
GO:0019842
vitamin binding
1.000000
7 / 268
GO:0043177
organic acid binding
1.000000
3 / 58
GO:0016984
ribulose−bisphosphat...
6.46e−05
5 / 24
GO:0030597
RNA glycosylase acti...
1.000000
4 / 26
GO:0140102
catalytic activity, ...
1.000000
4 / 95
GO:0043891
glyceraldehyde−3−pho...
1.000000
2 / 5
GO:0005542
folic acid binding
0.000993
2 / 3
GO:0030598
rRNA N−glycosylase a...
0.001209
4 / 26
GO:0004365
glyceraldehyde−3−pho...
0.003230
2 / 5

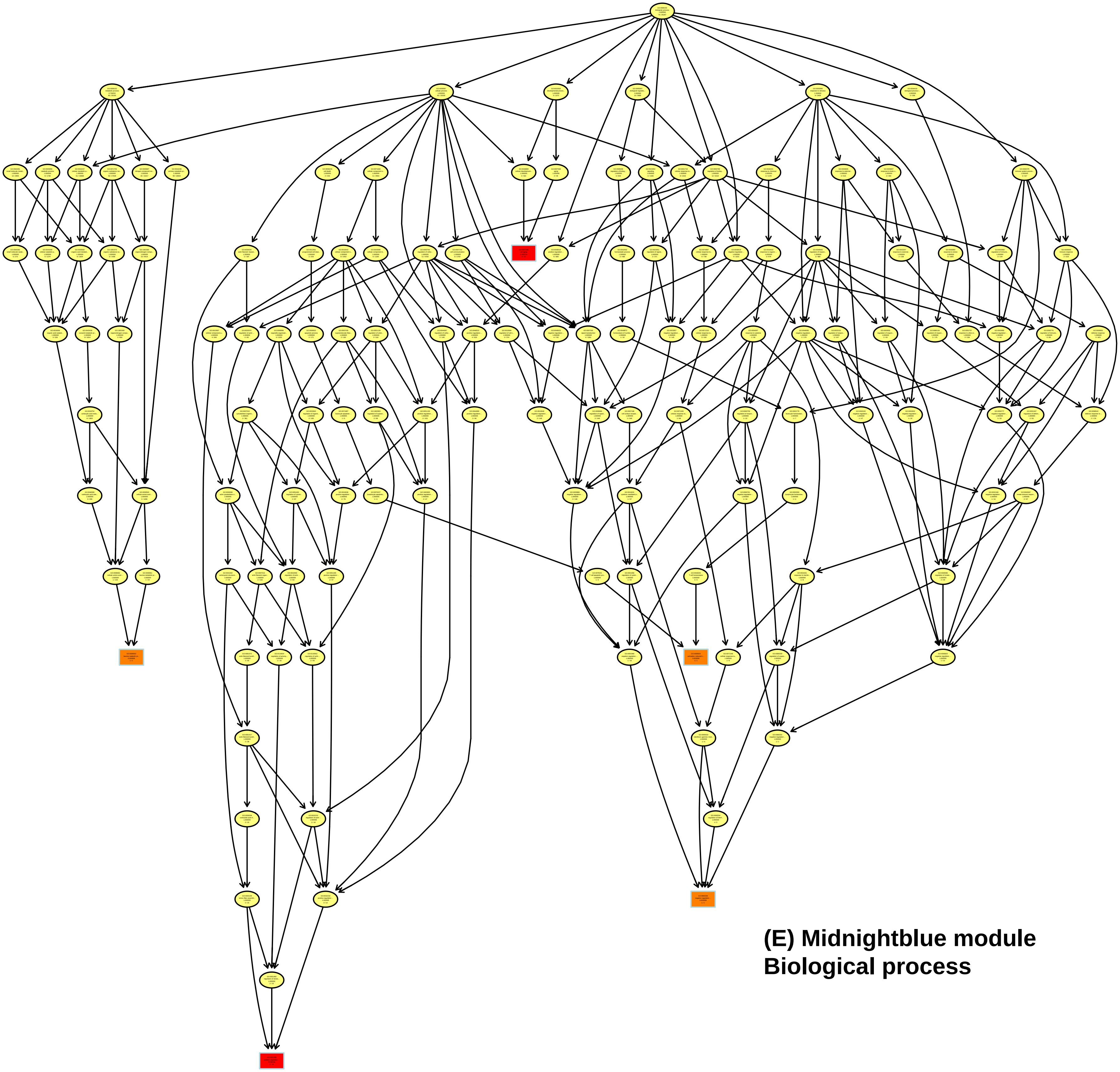

**(E) Midnightblue module Biological process**

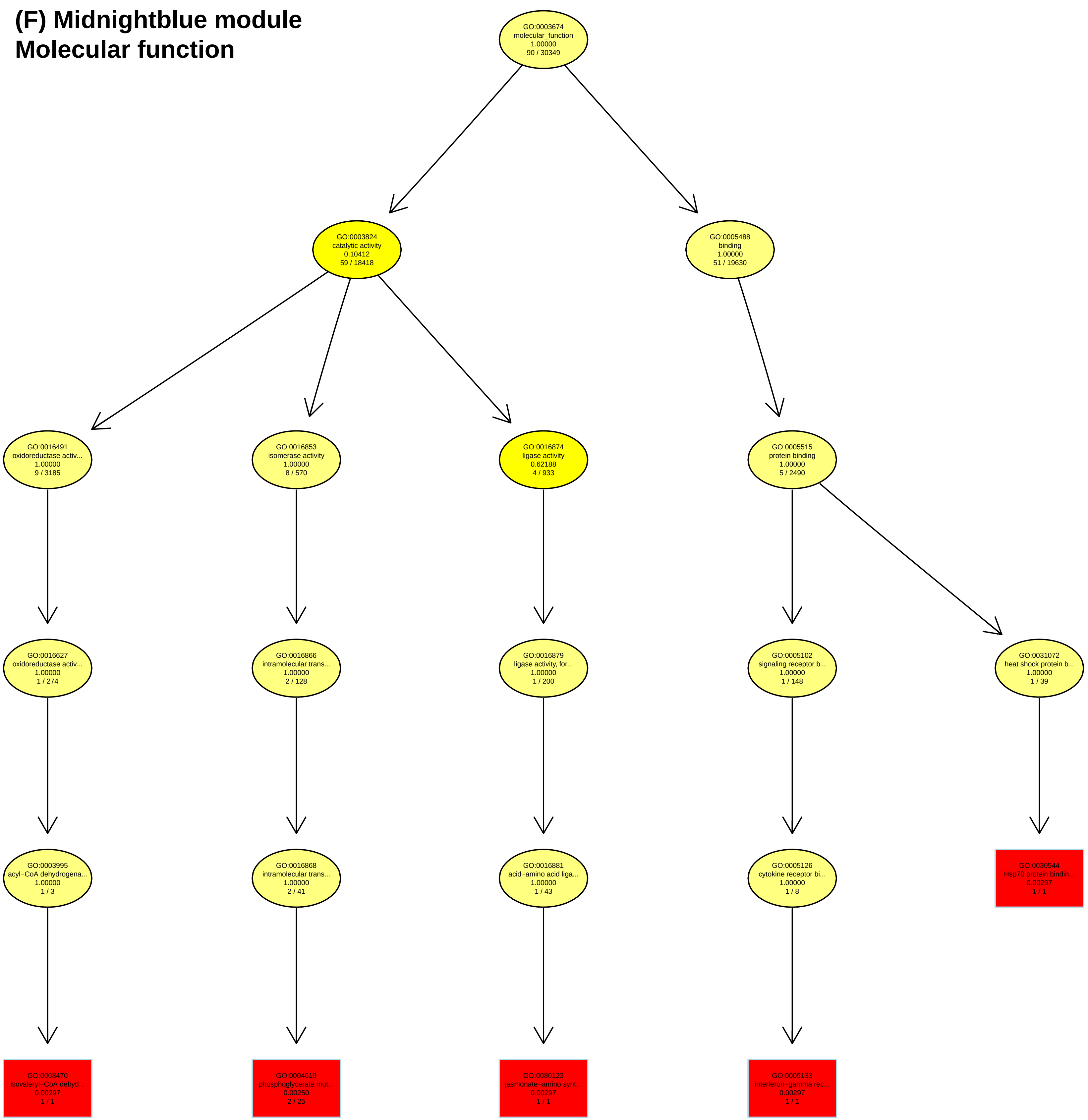
(F) Midnightblue module
Molecular function
GO:0003674 molecular_function 1.00000 90 / 30349
GO:0003824 catalytic activity 0.10412 59 / 18418
GO:0005488 binding 1.00000 51 / 19630
GO:0016491 oxidoreductase activ... 1.00000 9 / 3185
GO:0016853 isomerase activity 1.00000 8 / 570
GO:0016874 ligase activity 0.62188 4 / 933
GO:0005515 protein binding 1.00000 5 / 2490
GO:0016627 oxidoreductase activ... 1.00000 1 / 274
GO:0016866 intramolecular trans... 1.00000 2 / 128
GO:0016879 ligase activity, for... 1.00000 1 / 200
GO:0005102 signaling receptor b... 1.00000 1 / 148
GO:0031072 heat shock protein b... 1.00000 1 / 39
GO:0003995 acyl−CoA dehydrogena... 1.00000 1 / 3
GO:0016868 intramolecular trans... 1.00000 2 / 41
GO:0016881 acid−amino acid liga... 1.00000 1 / 43
GO:0005126 cytokine receptor bi... 1.00000 1 / 8
GO:0030544 Hsp70 protein bindin... 0.00297 1 / 1
GO:0008470 isovaleryl−CoA dehyd... 0.00297 1 / 1
GO:0004619 phosphoglycerate mut... 0.00250 2 / 25
GO:0080123 jasmonate−amino synt... 0.00297 1 / 1
GO:0005133 interferon−gamma rec... 0.00297 1 / 1

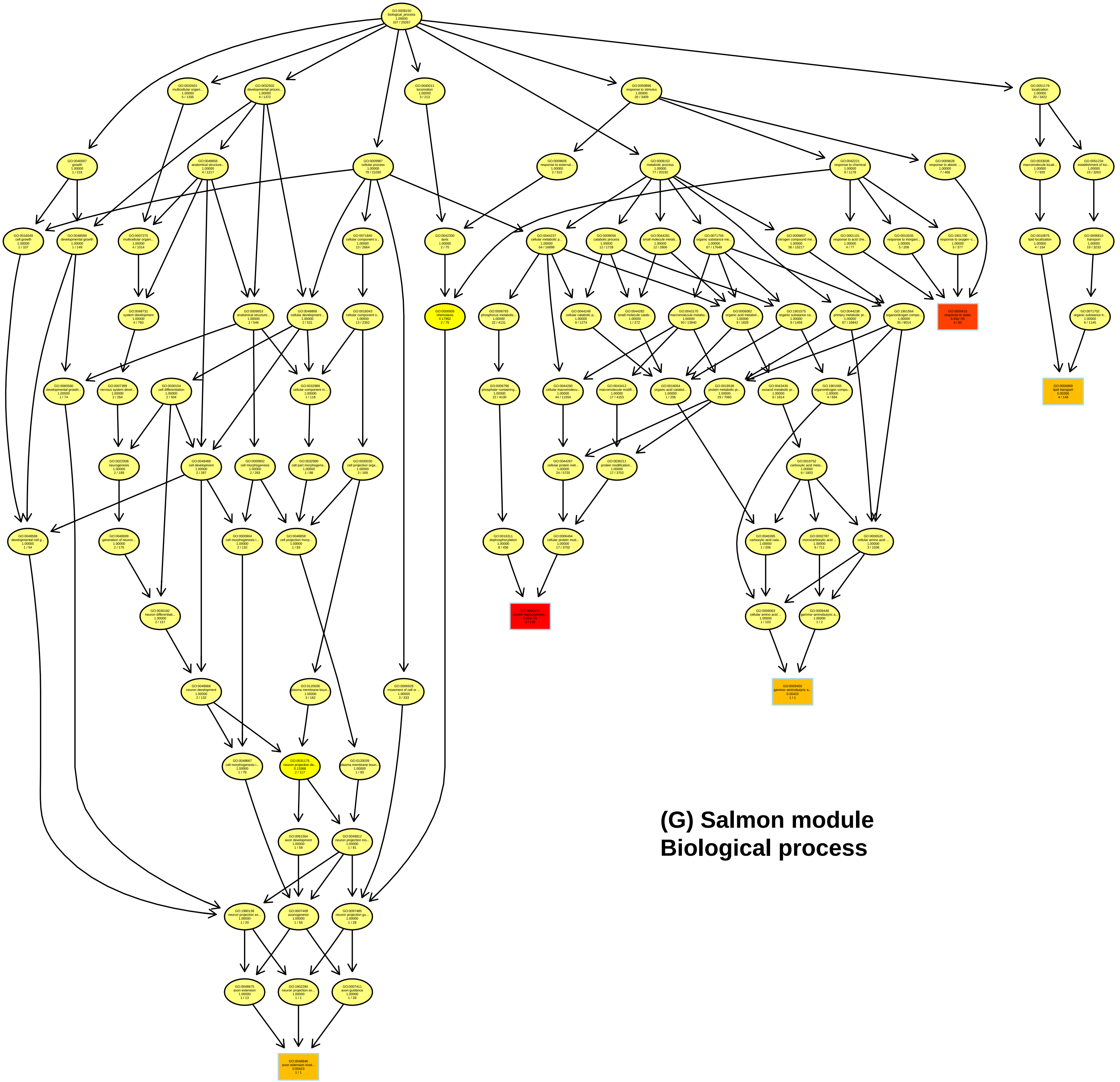

**(G) Salmon module**
**Biological process**

### (H) Salmon module
### Molecular function

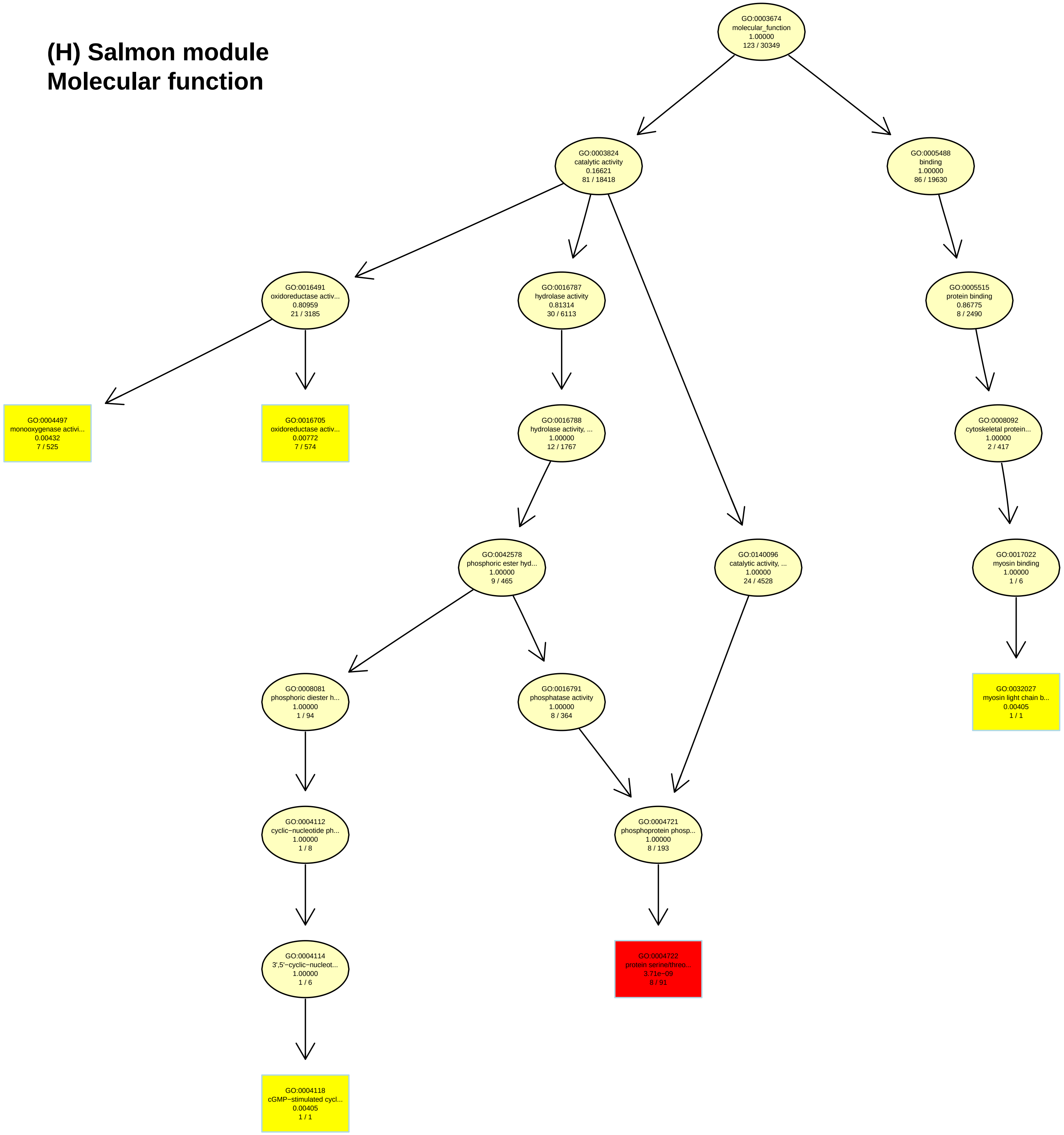

**(I) Turquoise module**
**Biological process**

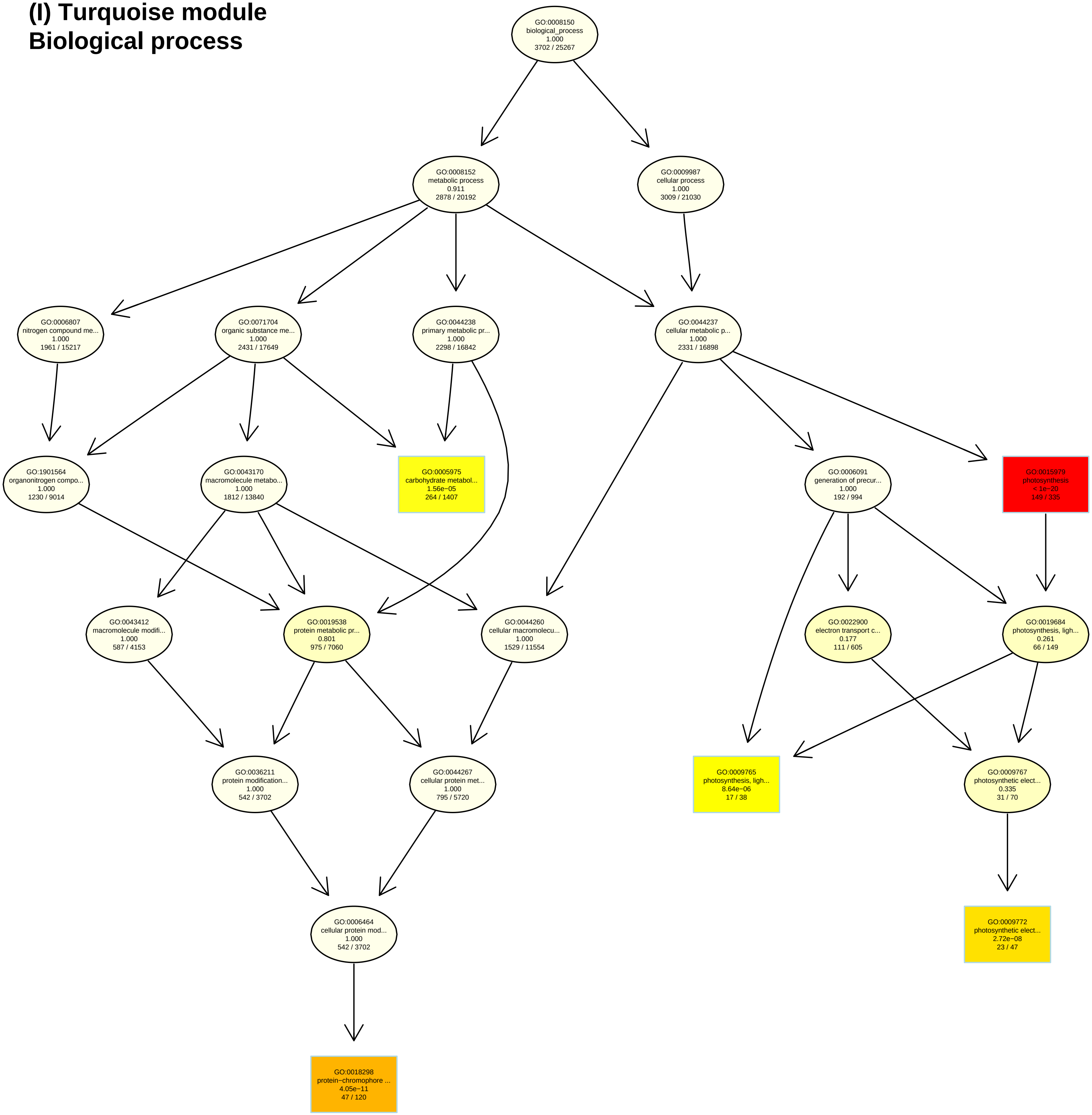

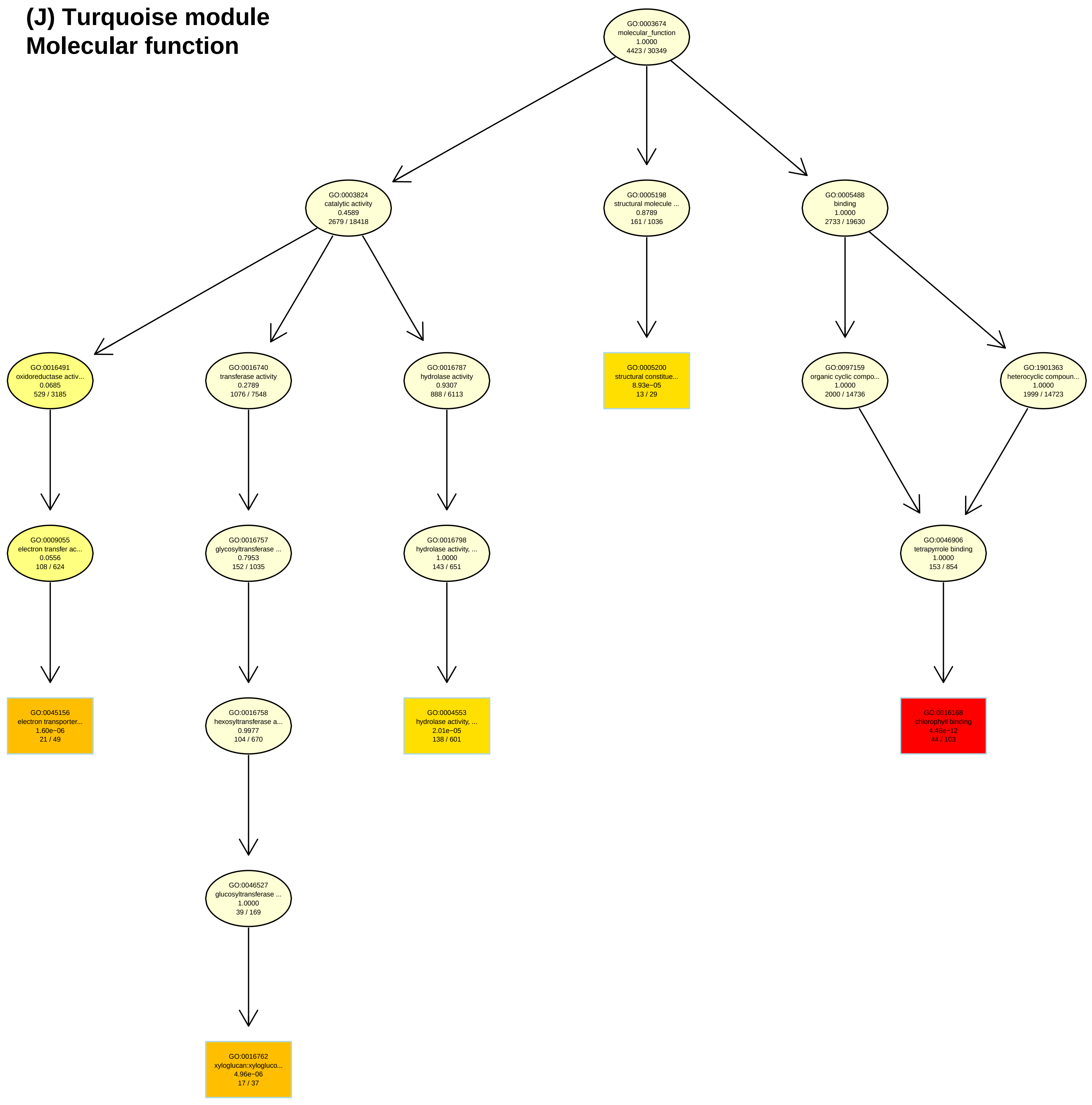
(J) Turquoise module
Molecular function
GO:0003674
molecular_function
1.0000
4423 / 30349
GO:0003824
catalytic activity
0.4589
2679 / 18418
GO:0005198
structural molecule ...
0.8789
161 / 1036
GO:0005488
binding
1.0000
2733 / 19630
GO:0016491
oxidoreductase activ...
0.0685
529 / 3185
GO:0016740
transferase activity
0.2789
1076 / 7548
GO:0016787
hydrolase activity
0.9307
888 / 6113
GO:0005200
structural constitue...
8.93e−05
13 / 29
GO:0097159
organic cyclic compo...
1.0000
2000 / 14736
GO:1901363
heterocyclic compoun...
1.0000
1999 / 14723
GO:0009055
electron transfer ac...
0.0556
108 / 624
GO:0016757
glycosyltransferase ...
0.7953
152 / 1035
GO:0016798
hydrolase activity, ...
1.0000
143 / 651
GO:0046906
tetrapyrrole binding
1.0000
153 / 854
GO:0045156
electron transporter...
1.60e−06
21 / 49
GO:0016758
hexosyltransferase a...
0.9977
104 / 670
GO:0004553
hydrolase activity, ...
2.01e−05
138 / 601
GO:0016168
chlorophyll binding
4.48e−12
44 / 103
GO:0046527
glucosyltransferase ...
1.0000
39 / 169
GO:0016762
xyloglucan:xylogluco...
4.96e−06
17 / 37

**Hierarchical trees for gene ontology enrichment analyses.** Hierarchical trees for the 5 highest ranked GO terms for all genes clustered in the blue, green, midnightblue, salmon or turquoise module, respectively, for both biological process and molecular function.
**(A)** Blue module, Biological process
**(B)** Blue module, Molecular function
**(C)** Green module, Biological process
**(D)** Green module, Molecular function
**(E)** Midnightblue module, Biological process
**(F)** Midnightblue module, Molecular function
**(G)** Salmon module, Biological process
**(H)** Salmon module, Molecular function
**(I)** Turquoise module, Biological process
**(J)** Turquoise module, Molecular function
