## Supplementary material for "Barley shows reduced Fusarium Head Blight under drought and modular expression of differential expressed genes under combined stress": suppl.FiguresS1-S3

### Supplemental figures

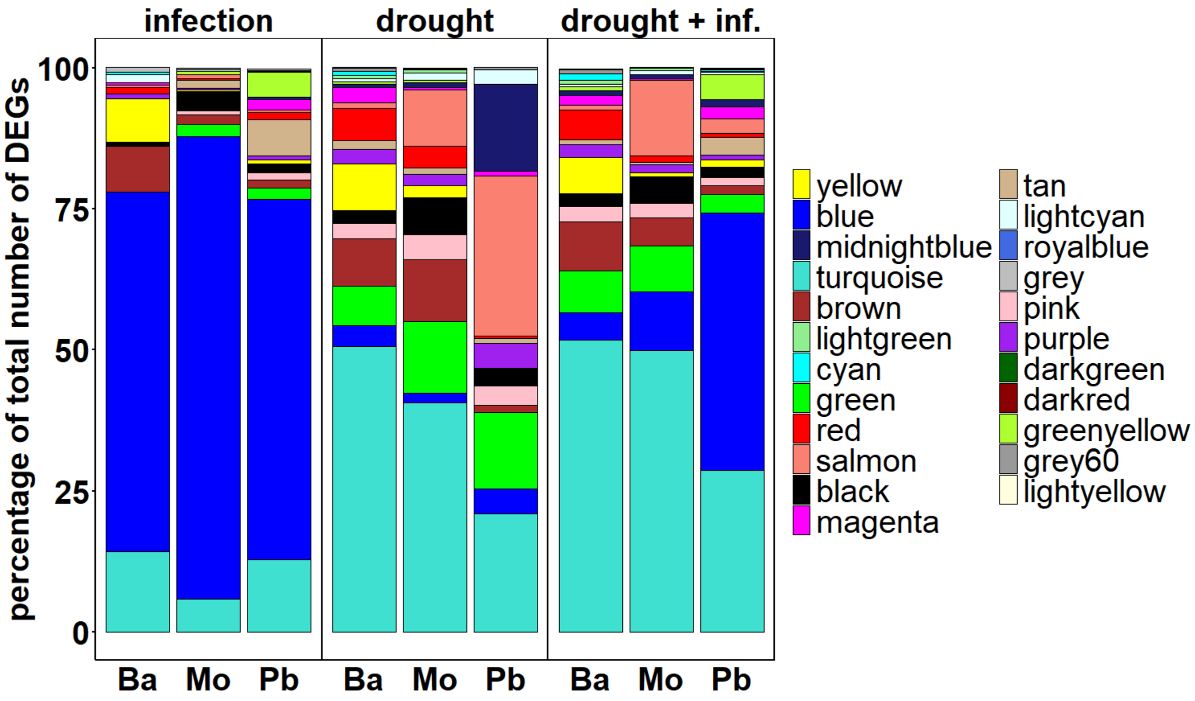


**Suppl. Fig. S1.** Full list of co-expression modules in each variety for each stress treatment. Number of DEGs was counted in each module for each variety under each stress treatment. The figure shows all modules for each scenario in percentage relative to the total number of DEGs per variety per treatment.

**A**

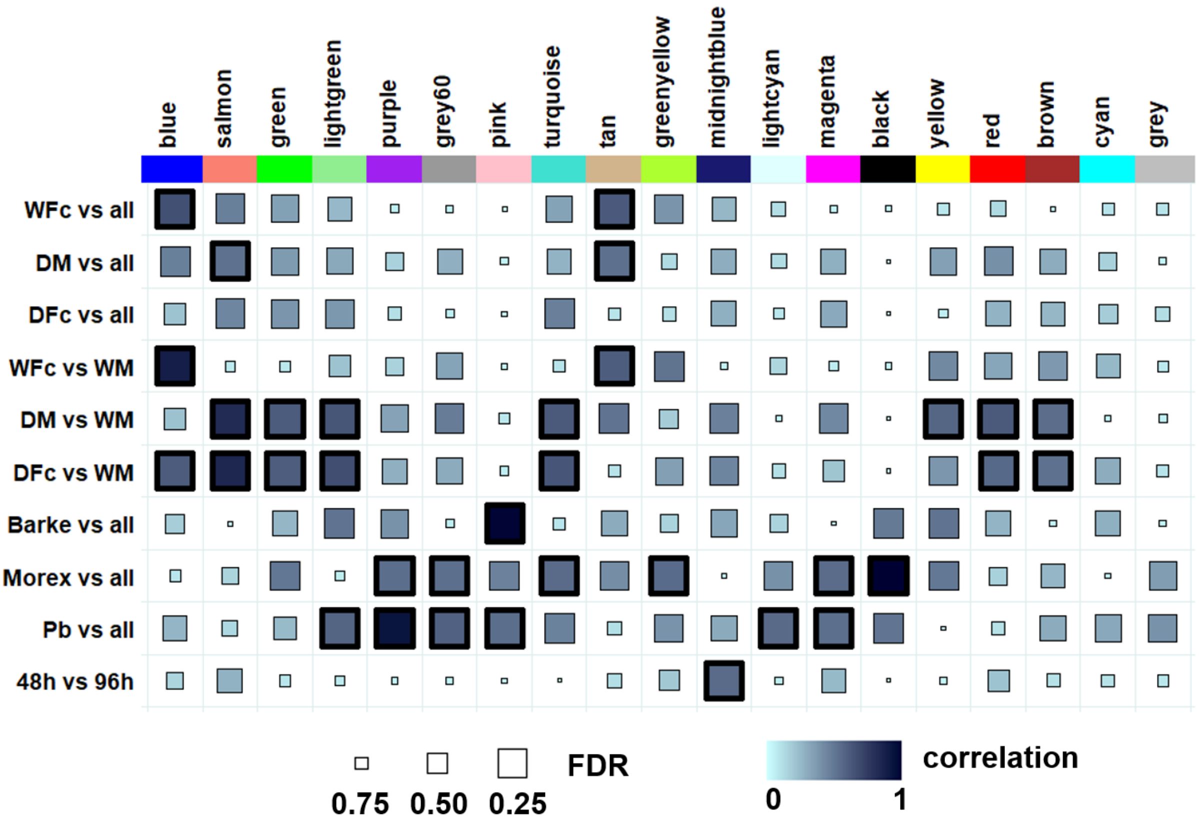


**B**

| | Barke vs all | Morex vs all | Pb vs all | DFc vs WM | DM vs WM | WFc vs WM | DFc vs all | DM vs all | WFc vs all | 48h vs 96h |
|---|---|---|---|---|---|---|---|---|---|---|
| Barke_D_Fc_48 | 1 | 0 | 0 | 0 | NA | NA | 1 | 0 | 0 | 0 |
| Barke_D_Fc_96 | 1 | 0 | 0 | 0 | NA | NA | 1 | 0 | 0 | 1 |
| Barke_D_Mock_48 | 1 | 0 | 0 | NA | 0 | NA | 0 | 1 | 0 | 0 |
| Barke_D_Mock_96 | 1 | 0 | 0 | NA | 0 | NA | 0 | 1 | 0 | 1 |
| Barke_W_Fc_48 | 1 | 0 | 0 | NA | NA | 0 | 0 | 0 | 1 | 0 |
| Barke_W_Fc_96 | 1 | 0 | 0 | NA | NA | 0 | 0 | 0 | 1 | 1 |
| Barke_W_Mock_48 | 1 | 0 | 0 | 1 | 1 | 1 | 0 | 0 | 0 | 0 |
| Barke_W_Mock_96 | 1 | 0 | 0 | 1 | 1 | 1 | 0 | 0 | 0 | 1 |
| Morex_D_Fc_48 | 0 | 1 | 0 | 0 | NA | NA | 1 | 0 | 0 | 0 |
| Morex_D_Fc_96 | 0 | 1 | 0 | 0 | NA | NA | 1 | 0 | 0 | 1 |
| Morex_D_Mock_48 | 0 | 1 | 0 | NA | 0 | NA | 0 | 1 | 0 | 0 |
| Morex_D_Mock_96 | 0 | 1 | 0 | NA | 0 | NA | 0 | 1 | 0 | 1 |
| Morex_W_Fc_48 | 0 | 1 | 0 | NA | NA | 0 | 0 | 0 | 1 | 0 |
| Morex_W_Fc_96 | 0 | 1 | 0 | NA | NA | 0 | 0 | 0 | 1 | 1 |
| Morex_W_Mock_48 | 0 | 1 | 0 | 1 | 1 | 1 | 0 | 0 | 0 | 0 |
| Morex_W_Mock_96 | 0 | 1 | 0 | 1 | 1 | 1 | 0 | 0 | 0 | 1 |
| Palmella_Blue_D_Fc_48 | 0 | 0 | 1 | 0 | NA | NA | 1 | 0 | 0 | 0 |
| Palmella_Blue_D_Fc_96 | 0 | 0 | 1 | 0 | NA | NA | 1 | 0 | 0 | 1 |
| Palmella_Blue_D_Mock_48 | 0 | 0 | 1 | NA | 0 | NA | 0 | 1 | 0 | 0 |
| Palmella_Blue_D_Mock_96 | 0 | 0 | 1 | NA | 0 | NA | 0 | 1 | 0 | 1 |
| Palmella_Blue_W_Fc_48 | 0 | 0 | 1 | NA | NA | 0 | 0 | 0 | 1 | 0 |
| Palmella_Blue_W_Fc_96 | 0 | 0 | 1 | NA | NA | 0 | 0 | 0 | 1 | 1 |
| Palmella_Blue_W_Mock_48 | 0 | 0 | 1 | 1 | 1 | 1 | 0 | 0 | 0 | 0 |
| Palmella_Blue_W_Mock_96 | 0 | 0 | 1 | 1 | 1 | 1 | 0 | 0 | 0 | 1 |

**Suppl. Fig. S2.** Relationships of consensus module eigengenes with binary comparisons. Each column in the table (A) corresponds to a module and each row to one of the indicated, binary comparisons shown in (B). (A) Module names are shown on top. Square colours in the figure represent the correlations of corresponding module eigengenes and defined comparison. The negative FDR-corrected p-values are coded by size (the bigger the square, the more significant). Highly significant correlations ($p < 0.01$) are highlighted with bold frames. (B) Listed in the table are all samples used in this study and their corresponding set values for each comparison.

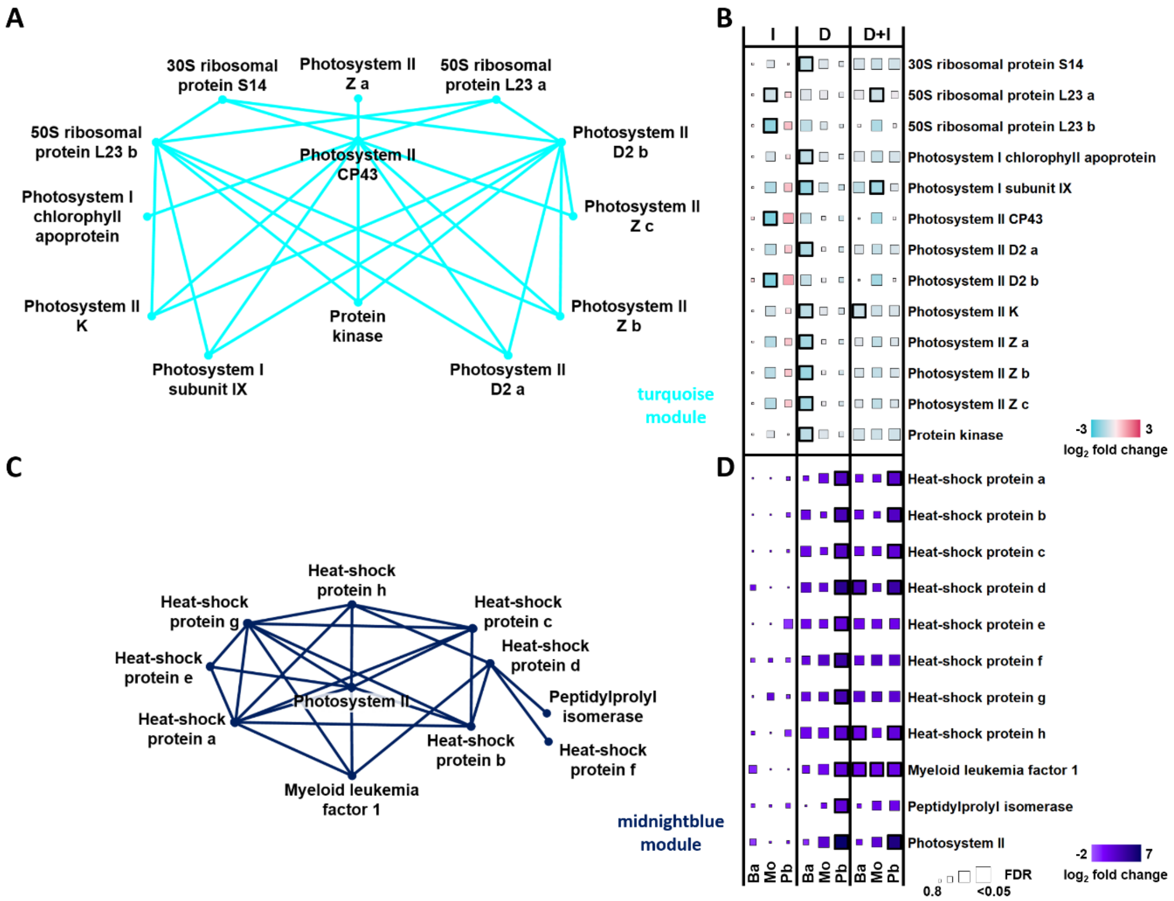


**Suppl. Fig. S3.** Co-expression levels reveal potential hub genes in selected modules. For two additional modules, midnightblue and turquoise, the 25 edges with the highest edge weight have been selected and their corresponding network was schematically drawn (A, C). Each node represents one gene and in the heat maps the log fold changes of the gene expression values were shown with the respective colour codes and individual scales for each module (B, D). The size of the square represents the false discovery rate (FDR), with bigger squares indicating higher confidence levels. The bold squares indicate a significant FDR p-value smaller than 0.05. For gene identifiers, see supplemental dataset D5.
