## Supplementary material for "Barley shows reduced Fusarium Head Blight under drought and modular expression of differential expressed genes under combined stress": suppl.T1-variety-information

**Suppl. table T1.** Detailed information about the three used barley varieties.

| **cultivar** | **row type** | **maturing type** | **traits** | **type** | **improvement status** | **pedigree** | **year of release** | **breeder** | **origin** | **disease resistance** | **source** |
| --- | --- | --- | --- | --- | --- | --- | --- | --- | --- | --- | --- |
| Barke | 2 | late | high malting properties, high yield | spring barley | modern cultivar | Libelle x Alexis | 1996 | Saatzucht Josef Breun GmbH & Co. KG | Southern Germany | resistant against powdery mildew (mlo9), quant. brown rust resistance | (Friedt *et al.* 2011) |
| Morex | 6 | late | high fermentable extract; barley reference genome | spring barley | cultivar | Cree x Bonanza | 1978 | Dr. Donald Rasmusson - University of Minnesota | Northern America | quant. resistance against spot blotch, stem rust,  bacterial leaf streak; susceptible towards net blotch | (Rasmusson and Wilcoxson 1979; Steffenson *et al.* 1996; El Attari *et al.* 1998) |
| Palmella Blue | 2 | early | dwarf, little lodging, drought tolerance | spring barley | landrace | unknown | unknown | unknown | Ethiopia | low disease resistance,  susceptible to Fusarium head blight | (Harlan H. *et al.* 1940; Lanzinger *et al.* 2015; Pourkheirandish *et al.* 2018; Hoheneder *et al.* 2022) |
